# The restricted N-glycome of neurons is programmed during differentiation

**DOI:** 10.1101/2024.10.15.618477

**Authors:** Katherine Kiwimagi, Maxence Noel, Murat Cetinbas, Ruslan I. Sadreyev, Lei Wang, Jordan W. Smoller, Richard D. Cummings, Ron Weiss, Robert G. Mealer

## Abstract

The protein glycome of individual cell types in the brain is unexplored, despite the critical function of these modifications in development and disease. In aggregate, the most abundant asparagine (N-) linked glycans in the adult brain are high mannose structures, and specifically Man_5_GlcNAc_2_ (Man-5), which normally exits the ER for further processing in the Golgi. Mannose structures are uncommon in other organs and often overlooked or excluded in most studies. To understand cell-specific contributions to the unique brain N-glycome and its abundance of Man-5, we performed RNAseq and MALDI-MS TOF protein N-glycomics at several timepoints during differentiation of multiple cell types. To this end, homogeneous cultures of glutamatergic neurons, GABAergic neurons, and brain-specific endothelial cells were generated from monoclonal human inducible pluripotent stem cells (hiPSCs) through cellular reprogramming. Small molecule induction of stably integrated synthetic transcription units driving morphogen expression generated differentiated cells with distinct patterns mirroring intact tissue. Comparing uninduced hiPSCs for each cell type revealed identical transcriptomic and glycomic profiles before differentiation, with low quantities of Man-5. In differentiated glutamatergic and GABAergic neurons, the most abundant N-glycans became Man-5 and its immediate precursor Man-6, despite the presence of transcripts encoding enzymes for their subsequent modification. Differentiation to brain-specific endothelial cells showed an opposite effect, with the N-glycome displaying an abundance of complex N-glycans and terminal modifications of the late secretory pathway. These results confirm that the restricted N-glycome profile of brain is programmed into neuronal differentiation, with regulation independent of the transcriptome and under tight evolutionary constraint.

## Introduction

N-glycosylation is an evolutionarily conserved biological pathway involving the enzymatic attachment of carbohydrate polymers to asparagine residues of proteins. N-glycosylation is critical in the development and function of the brain, as congenital disorders associated with the pathway most commonly present with severe neurologic phenotypes including seizures and intellectual disability^1^. Growing evidence is also revealing that common genetic variants in glycosylation genes are associated with more complex neuropsychiatric phenotypes including schizophrenia and Alzheimer’s disease^2,3^. One such example is mutations in the manganese (Mn) transporter *SLC39A8*, as many glycosylation enzymes require Mn as a co-factor for activity. Severe mutations in *SLC39A8* result in a type II congenital disorder of glycosylation characterized by near total loss of circulating Mn and profound psychomotor impairment, epilepsy, and growth abnormalities^4,5^. On the opposite end of the allelic spectrum, a common missense variant (Ala391Thr - A391T, rs13107325) in *SLC39A8* is associated with hundreds of complex human phenotypes by GWAS^6^, including decreased serum Mn^7^, schizophrenia^8^, intelligence^9^, and several neuroimaging findings^10,11^. Both human carriers and mouse models of A391T have slightly reduced circulating Mn and abnormal serum glycosylation^12–14^, and the A391T mouse model has altered protein glycosylation in the brain^15^. Interestingly, in the brain, *SLC39A8* is almost exclusively expressed in endothelial cells and absent in neurons^16^.

Protein N-glycosylation is present in all domains of life but thought to be most complex in mammals^17^. All eukaryotic cells initiate N-glycosylation on the cytoplasmic side of the ER, where a dolichol-phosphate anchor is elongated with monosaccharides and then flipped into the lumen. This lipid-oligosaccharide complex is further extended in the ER prior to transfer onto asparagines (N-linked) of recently translated proteins entering the secretory pathway. The N-glycan precursor is then trimmed back by a series of α-glucosidases to generate the structure Man_9_GlcNAc_2_ (Man-9) and Man_8_GlcNAc_2_ (Man-8), which are common structures on glycoproteins exiting the ER. Mannose residues are then sequentially cleaved in the cis-Golgi by a series of α-mannosidases to generate Man_5_GlcNAc_2_ (Man-5). After high mannose (Man-5 through Man-9) structures, generation of complex N-glycans in all cells requires the action of the N-acetylglucosaminyltransferase I (MGAT1), which adds a GlcNAc residue to Man-5. MGAT1 is an essential enzyme, broadly expressed across all tissues with knock-out mice dying around embryonic day 10 and showing notable impairments in neural development^18^. After MGAT1, N-glycans can be modified by α-mannosidase II and hundreds of different glycosyltransferases and related enzymes to generate a seemingly infinite array of glycan structures.

Most tissues and plasma display an abundance of complex N-glycans with relatively few high mannose structures, generally less than <20%^19,20^. Some studies even exclude high mannose structures from analysis, presumably as precursors for more complex and biologically relevant glycans^21^. The brain N-glycome, however, is predominantly high mannose structures by bulk (>60%), with Man-5 being the most abundant^22^. This observation is consistent across mammals^22,23^ as well as zebrafish^24^. We supposed one contributor to the unique N-glycome of the brain was broad transcriptional downregulation of glycosylation genes compared to other tissues^22^. In neurons, other groups have described unconventional secretory processing^25^, Golgi-independent trafficking^26^ and activity-dependent Golgi satellites^27^ as mechanisms which contribute to the unique qualities and function of N-glycans in the brain. However, the contribution of individual cell types as well as their developmental timeline responsible for the distinct brain N-glycome is unknown.

To help fill these gaps in understanding, we generated human iPSCs with or without the *SLC39A8* gene and subsequently differentiated them into glutamatergic neurons, GABAergic neurons, and endothelial cells using inducible gene circuits, expecting only endothelial cells to be affected by *SLC39A8* deletion. We confirmed specific differentiation through RNAseq analysis and used immunofluorescence and qPCR as additional probes of cell state when needed. MALDI-TOF MS revealed unique N-glycome profiles for both neuronal cell types and endothelial cells, mirroring previous studies of intact tissue and primary cells^22,28^. Genomic deletion of *SLC39A8* had minimal effect on the N-glycome of glutamatergic and GABAergic neurons, whereas loss of this gene in endothelial cells where it is normally expressed caused a dramatic reduction of complex N-glycans. These findings are the first detailed glycosylation studies of individual human brain cell types and highlight the distinct and complex regulation of this critical modification in the nervous system.

## Results

### Morphogen expression via stably integrated inducible circuits drives targeted cellular differentiation

To study the effect of differentiation on the N-glycome of distinct cell types, gene driver circuits were stably integrated into human induced pluripotent stem cells (hiPSCs) with a shared isogenic background (PGP1)^29^. One endothelial and two neuronal cell types were targeted via doxycycline (DOX) induced morphogen expression in gene driver circuits with previously established differentiation protocols optimized for generation of homogeneous monocultures free of support cells (**Fig. 1A**). The targeted neuronal cell types were based on inducible expression of *<u>N</u>GN1* and *<u>N</u>GN2* (iNN) for glutamatergic neurons^30^ and inducible expression of *<u>A</u>SCL1* and *<u>D</u>LX2* (iAD) for GABAergic neurons^31^. In a similar fashion, brain-specific endothelial cell induction was based on inducible expression of *<u>E</u>TV2* (iE)^32^, though there is some controversy as to whether *ETV2* expression alone is sufficient to achieve a brain specific endothelial phenotype^33^. This controversy may be due to gaps in our knowledge on the effect of multiple transcription factors and the interplay between them, as well experimental variables such as varying growth media using different markers to define cell type. We therefore spent additional time characterizing resultant endothelial cells to confirm their transcriptomic and functional state. Circuits were stably integrated using Lenti or Piggybac into PGP1s followed by monoclonal sorting as described in Methods. After stable integration and gated sorting, monoclonal cell lines containing gene driver circuits were selected for differentiation into homogeneous populations through DOX induction (+DOX), with subsequent analyses performed on days 0, 4, and 14 (**Fig. 1B).**

**Figure 1.**
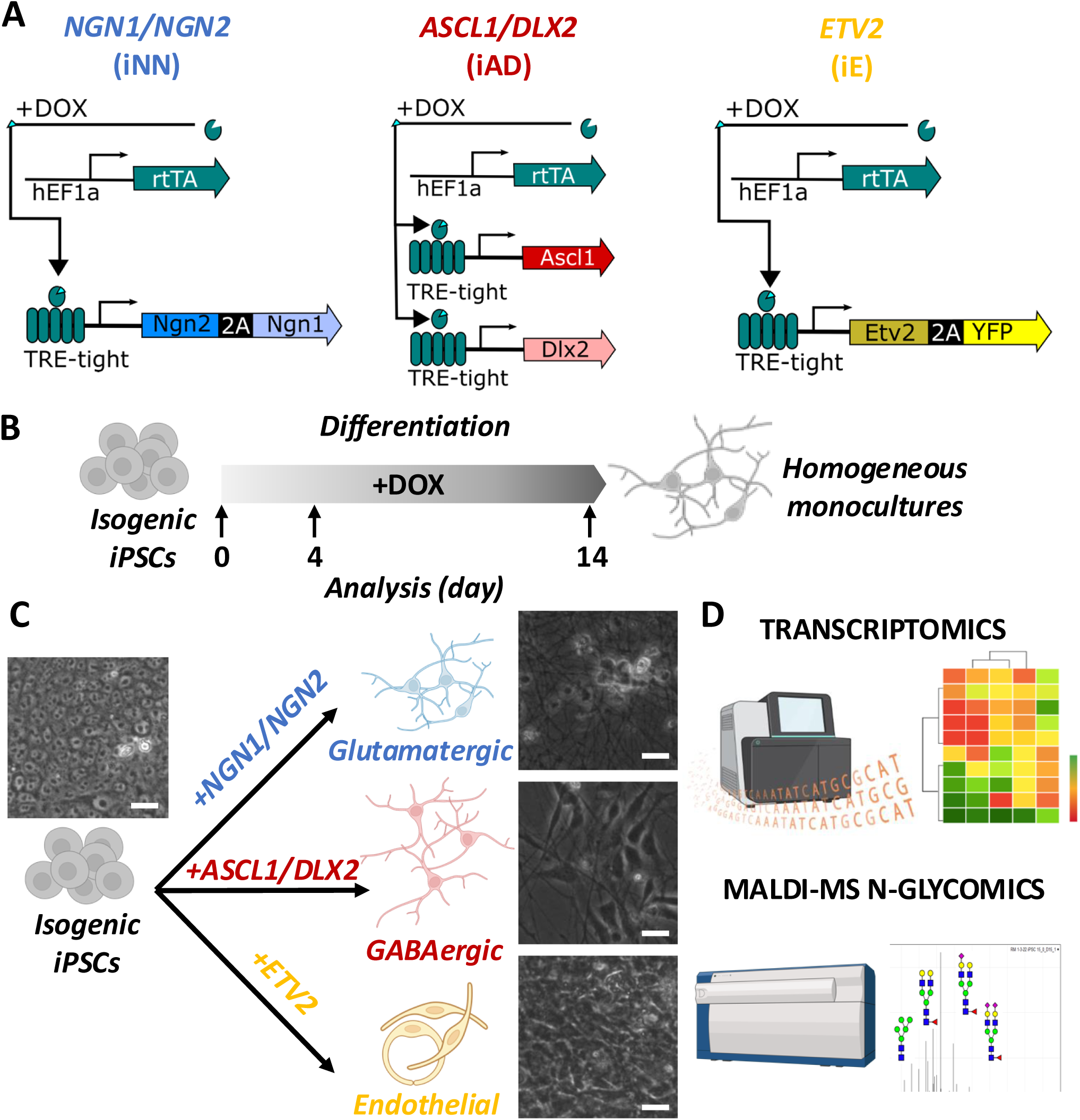
Defining the N-glycome of single human cell types. **A)** Schematic of differentiation and analysis of Isogenic human iPSCs used to generate homogeneous monocultures of distinct cell types following doxycycline (+DOX) induction. **B)** Gene circuit designs for differentiation of glutamatergic neurons (iNN), GABAergic neurons (iAD), and endothelial cells (iE) based on established protocols. **C)** Schematic and light microscopy images showing differentiation of iPSCs into homogeneous monocultures with morphology reflecting distinct cell types. Scale bar = 20 µm. Uncropped images are presented in supplemental material. **D)** Analytic methods including RNAseq and MALDI-TOF MS N-glycomics applied at different time points confirm cell-specific transcriptome and glycome profiles.

In addition, a pair of otherwise identical cells lacking the expression of the Mn transporter SLC39A8 (*SLC39A8-/-*) was created using CRISPR/Cas9 and verified using genomic PCR for each gene driver set cell line (**Supp. Fig. 1**). As *SLC39A8* expression in the brain is limited exclusively to endothelial cells^16^, its deletion would be expected to have little to no effect on neuronal cell types in homogeneous cultures. This resulted in 3 isogenic pairs (6 cell lines total) to investigate N-glycome differences during differentiation. Transformation of hiPSCs to differentiated monocultures was first confirmed by cellular morphology (**Fig. 1C**), followed by in depth parallel analyses using bulk RNAseq transcriptomics and MALDI-TOF MS N-glycomics (**Fig. 1D**).

Human iPSCs grown in culture without DOX induction (-DOX) maintained an iPSC phenotype, with a patchy, sheet-like morphology and clusters of round cells seen on high magnification (**Supp. Fig. 2**). After 14 days +DOX, both glutamatergic iNN and GABAergic iAD neurons showed a less dense culture containing condensed somas and numerous neurite extensions. In contrast, iE endothelial cells generated a dense monolayer of elliptical-appearing cells.

Immunofluorescence for the iNN cell line with antibodies for the pluripotency markers SOX2 and OCT4 showed robust expression in the -DOX condition, which was then lost after 4 days +DOX (**Supp. Fig. 3**). Expression of NeuN was present in the cytosol of -DOX iNN cells, consistent with prior studies of undifferentiated cells (**Supp. Fig. 4**). After 4 days +DOX, expression of NeuN localized to the nucleus, consistent with a neuronal phenotype^34^. Expression of the neurofilament protein Tuj1 was absent in the -DOX iNN cells, but exhibited a strong signal in neuronal processes with 4 days +DOX. These results demonstrate on a morphological and protein level the conversion from iPSCs to differentiated cell types after induction.

### RNAseq confirms cell identity and differentiation state of homogeneous cultures

Bulk RNAseq was performed from each cell line before and after differentiation aligned to the human genome. Common cell markers for undifferentiated pluripotent cells (*DPPA4*, *NANOG*, *SOX2*), neurons (*MAP2*, *GRIA2*, *NCAM1*), glutamatergic neurons (*VGLUT1*, *SATB2*, *SYN1*), GABAergic neurons (*DLX1*, *GAD2*, *GAT1*), and endothelial cells (*FLI1*, *ERG*, *TAL1*) were compared across different cell types and showed enriched expression in each of the intended cell types (**Fig. 2**). Additional support for a brain-endothelial cell phenotype of ETV2 cell lines included the presence of mature endothelial cell markers *CDH5*, *CLDN5*, VWF, *FLT1*, and *TIE1,* and the reduction of epithelial markers *MUC1*, *EPCAM*, *CDH3*, and *CDH1* (**Supp. Table 1**).

**Figure 2.**
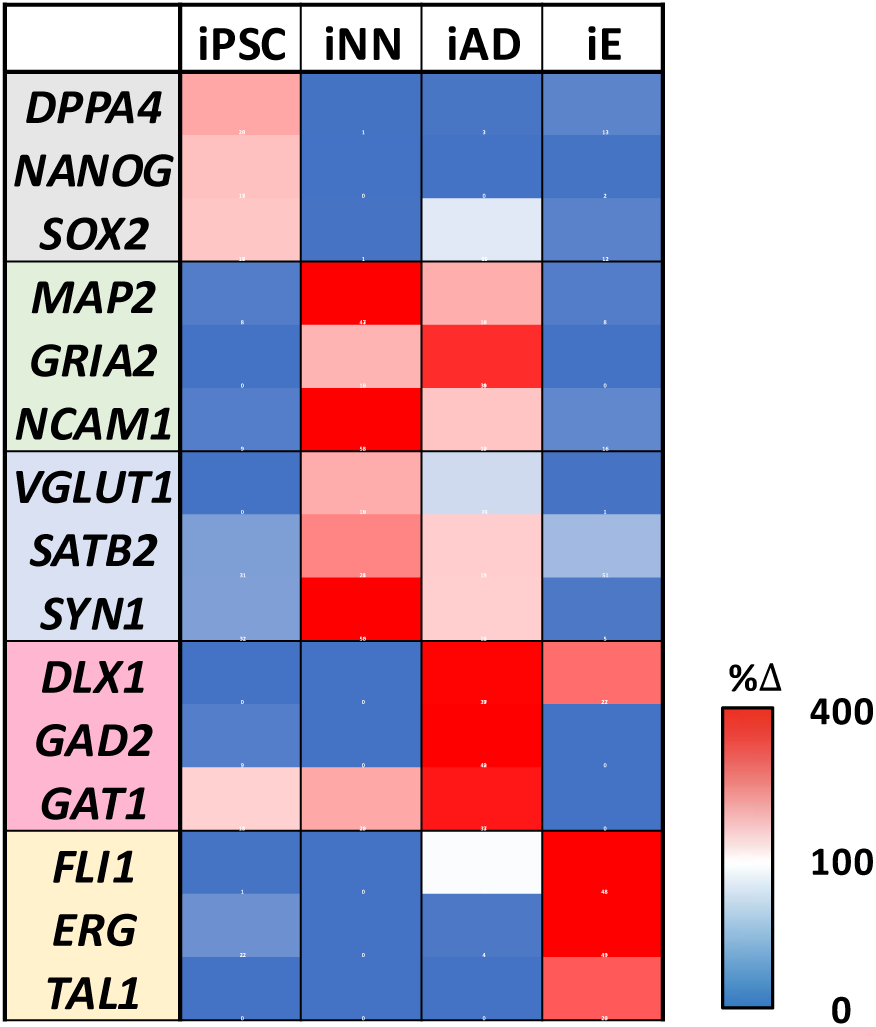
Transcriptomic profiles are consistent with programmed cell type and differentiation state. Relative expression of common cell-type markers are shown for iPSCs (gray), all neurons (green), glutamatergic neurons (blue), GABAergic neurons (red), and endothelial cells (yellow). Data is presented as percent change (%Δ) of individual cultures compared to average expression across all cultures. Heat map scaled from 0-400 % based on average expression across cultures.

RNAseq results for the induced morphogen followed the expected trend for iNN and iAD; however, in iE cells, the uninduced population exhibited unexpected high expression of *ETV2* at baseline (**Supp. Fig. 5**). To investigate this further, we performed qPCR using ETV2-specific primers on multiple +/- DOX iE samples, which confirmed low expression of *ETV2* at baseline in uninduced cells that increased in the presence of DOX (**Supp. Fig. 6**). As uninduced iE cell lines maintain the iPSC morphology and expression of iPSC markers such as *NANOG* and *SOX2,* we hypothesize that the confounding RNAseq results were a result of a lack in RNAseq probe sensitivity or an off-target artifact of the iE genetic engineering cassette, though it did not appear to affect differentiation or morphology.

A MDS plot comparing the global transcriptomic profile across all cell lines was consistent with specific and distinct cell types (**Fig. 3**). All 6 undifferentiated iPSCs, independent of the gene driver cassette, showed tight clustering of their transcriptomes. Following differentiation, iE lines were closely associated and distinct from the other clusters, as were the iNN and iAD neuronal cell lines. *SLC39A8* genotype appeared to have minimal effect on the transcriptomic profile of the cell lines in comparison to +/-DOX and the gene driver expression cassette, though iNN KO cells had minor separation from WT cells. These results are consistent with differentiation from pluripotency to homogeneous and mature cell cultures at the RNA level.

**Figure 3.**
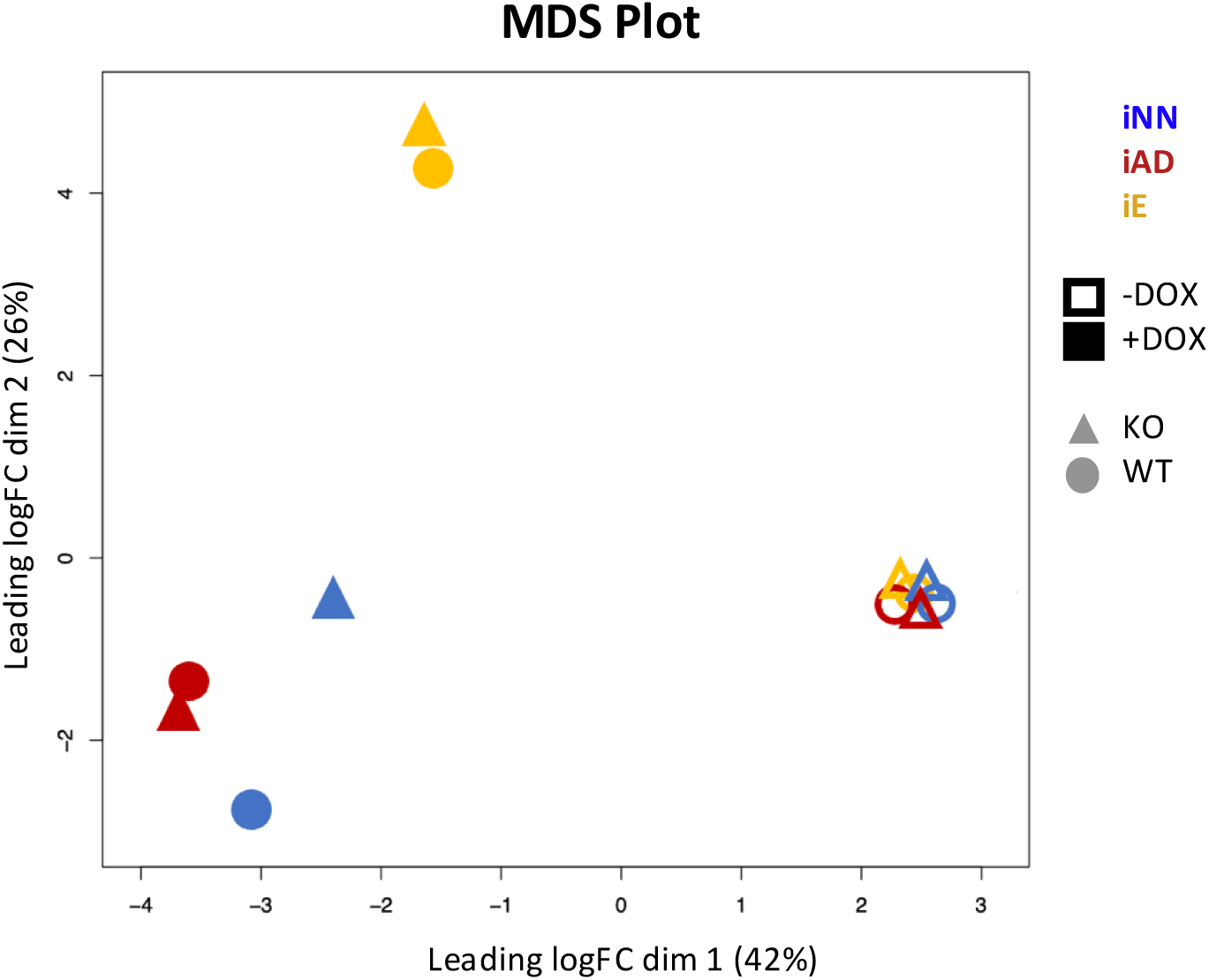
Transcriptomic profiles of distinct cell types form tight clusters. MDS plot of transcriptomic profiles from all cultures, highlighting the tight clustering of each undifferentiated iPSCs line (-DOX, open shapes) compared to induced cell lines (+DOX, filled shapes) expressing distinct gene drivers for iNN-neurons (blue), iAD-neurons (red) and iE-endothelial cells (yellow). Data is shown from two different genetic backgrounds, wild-type (circles) and *SLC39A8 -/*- (triangles) cells.

### The N-glycome is dynamic and cell-type specific during differentiation

We next performed MALDI-TOF MS glycomic analyzes of permethylated N-glycans prepared from each cell line at day 0, 4, and 14 of differentiation. For each set of cells, N-glycan masses were included if they had; 1- an isotopic appearance, 2 - a mass that corresponds to a possible glycan composition, and 3 - the average signal to noise ratio was greater than 6 (S/N>6). This resulted in the inclusion of 92, 80, 78, and 89 individual N-glycan masses for undifferentiated iPSCs, iNN, iAD, and iE cultures, respectively. Upon differentiation, unique N-glycome profiles emerged for each cell line, exhibiting distinctive overall profiles as well as exclusive peaks of highest abundance (**Fig. 4**, **Table 1**).

**Figure 4.**
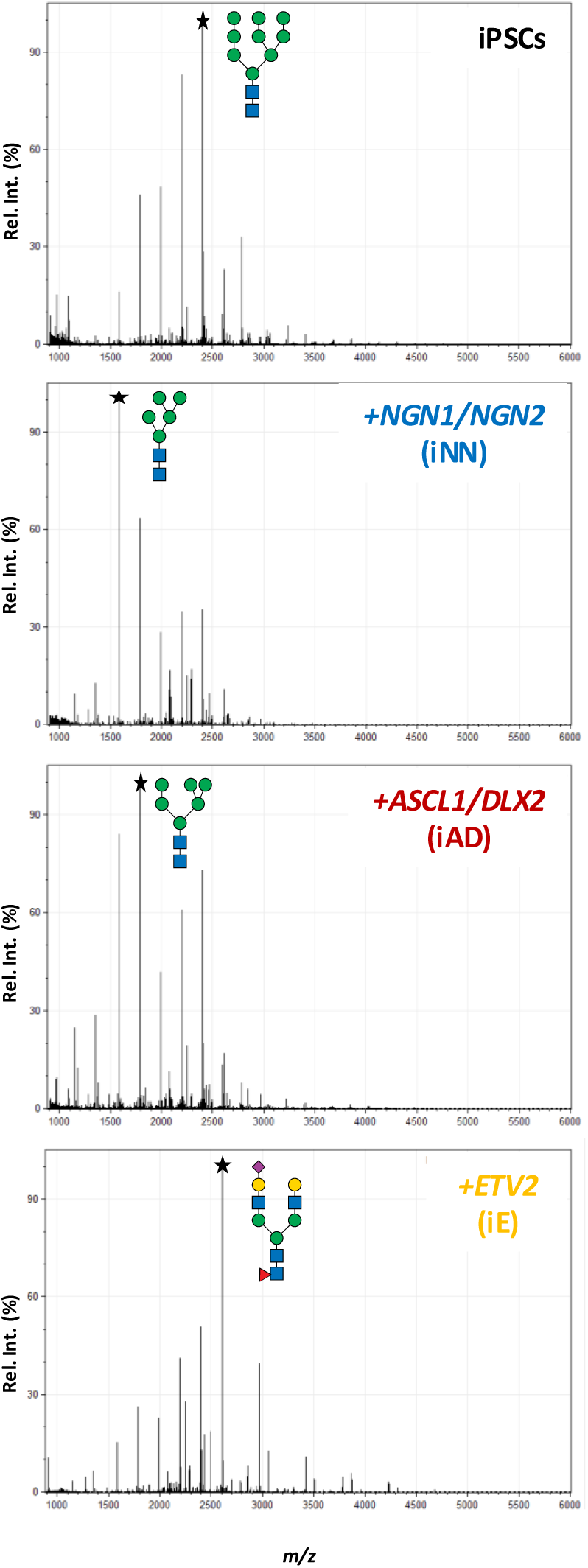
N-glycome profiles are unique between cell-type and resemble intact tissue. MALDI-MS TOF analysis of permethylated N-glycans isolated from homogeneous wild-type cultures of undifferentiated and differentiated iPSCs. The corresponding structure for the most intense peak (★) in each culture is illustrated, including Man-9 (iPSCs), Man-5 (iNN), Man-6 (iAD), and FA2G2S1 (iE).

**Table 1.** Glycan abundances vary between human cell types after 14 days of induced differentiation. Data is shown as the sum of the percent abundance for each category or glycan, normalized within the total N-glycome profile.

|  | **iPSC** | **iNN** | **iAD** | **iE** |
| --- | --- | --- | --- | --- |
| **Pauci-mannose** | 2.3 | 7.9 | 14.6 | 3.1 |
| **High-mannose** | 73.5 | 62.5 | 58.5 | 37.1 |
| **Complex** | 24.1 | 30.4 | 26.9 | 59.8 |
| **Sialic Acid** | 12.8 | 6.5 | 7.3 | 39.6 |
| **Fucose** | 21.2 | 27.0 | 29.6 | 52.6 |
| **Man-5** | 4.2 | 24.5 | 14.1 | 3.9 |
| **Man-6** | 12.8 | 15.6 | 16.7 | 6.7 |
| **Man-7** | 13.1 | 7.0 | 7.0 | 5.8 |
| **Man-8** | 21.7 | 7.8 | 9.5 | 9.8 |
| **Man-9** | 21.0 | 7.0 | 10.3 | 10.7 |

### The N-glycome of human iPSCs resemble other less differentiated cells

The N-glycome of undifferentiated iPSCs from every cell line was nearly identical (**Supp. Fig. 7**). High mannose structures composed ∼75% of the signal, with Man-9 and Man-8 being the most abundant structures, each independently representing ∼21% of the total signal (**Table 1**, **Supp. Fig. 8**). Of the complex N-glycans present (24% total), most contained a fucose (88%) and many were sialylated (53%). The N-glycome of human embryonic stem cells exhibited a similar pattern with high abundance of larger oligomannosidic structures including Man-8 and Man-9^35^. This pattern is also observed in studies of other cultured cells including HEK-293s^36^, HeLa^37^, and CHOs^38^ – with the exception of CHO cells lacking the N-acetylglucosaminyltransferase required for the final processing step of Man-5 to complex glycans, MGAT1, known as Lec1 cells^38^.

### The N-glycome of *NGN1/NGN2*-induced glutamatergic neurons (iNN) has abundant high mannose structures including Man-5

During *NGN1/NGN2* induction, iPSCs differentiating into glutamatergic lineage (iNN) exhibited a progressive shift to structures of smaller glycan masses by day 4 which continued through day 14 (**Supp. Fig. 9**). High mannose structures composed 62.5% of the signal, with Man-5 being the most abundant structure (24.5%) of the total signal, followed by Man-6 (15.6%) (**Table 1**, **Supp. Fig. 10**). Of the complex N-glycans present, a similar amount contained a fucose resideue compared to iPSCs, but far fewer were sialylated. Even the most abundant complex N-glycan detected in iNN cells exhibited apparent less abundance than any of the high mannose glycans, demonstrating the dramatic shift towards these glycans.

### The N-glycome of *ASCL1*/*DLX2*-induced GABAergic neurons (iAD) has abundant high mannose glycans including Man-6 and Man-5

After 4 days of *ASCL1*/*DLX2* induction, iPSCs differentiating into GABAergic lineage (iAD) showed a slower shift towards the structures that would be most abundant by day 14 (**Supp. Fig. 11**). At day 14, high mannose structures composed 58.5% of the signal, with Man-6 and Man-5 being the most abundant structures at 16.7% and 14.1% of the total signal, respectively (**Table 1**, **Supp. Fig. 12**). Of the complex N-glycans present, essentially all contained fucose, in addition to several pauci-mannose structures with fucose, and far fewer complex glycans contained sialic acid compared to both iNN and iPSCs. This pattern highlights a shift from complex and sialylated N-glycans to high mannose N-glycans during GABAergic neuron differentiation.

### The N-glycome of *ETV2*-induced endothelial cells (iE) displays robust sialylation and glycan complexity

At 4 days of *ETV2* induction, iPSCs differentiating into endothelial cells (iE) had begun to exhibit a reduction of high mannose glycans and an increase of more complex glycans (**Supp. Fig. 13**). At day 14, high mannose structures had decreased to 37.1 % of the signal, with Man-9 remained the most abundant structure of this class at 10.7% (**Table 1**, **Supp. Fig. 14**). Complex N-glycans had increased substantially to 59.8%, the majority of which contained both fucose and sialic acid. The complex biantennary glycan containing 2 galactoses, 1 fucose, and 1 sialic acid (FA2G2S1), had increased from 2.0% in iPSCs to 11.4% and 18.5% by day 4 and day 14, respectively, becoming the most abundant structure in iE cells (Supplementary Dataset 1). This data suggests that during differentiation to endothelial cells from iPSCs, the N-glycome transitions to more complex structures containing branches, fucose, and sialylation.

### The N-glycome of differentiated iPSCs tightly cluster and mirror intact tissue and primary cells

We next sought to quantitatively compare the glycomes of each cell line considering the entirety of the glycomic profile. Glycans with an isotopic appearance and mass that corresponds to known composition were selected at days 0, 4, and 14 of DOX induction for every sample. Individual *m/z* values present in all samples were extracted and normalized within each sample, followed by application of a Savitzky-Golay filter to reduce noise. A partial least squares regression (PLSR) against days of DOX induction resulted in an optimal number of 4 components, and plotting the first two components revealed clear groupings (**Fig. 5**). Resembling our RNAseq analysis, all 6 undifferentiated iPSCs showed tight clustering of their glycomes, consistent with their overall similar profiles (**Supp. Fig. 7**). After 4 days of DOX induction, each cell line had diverged from their parental iPSC glycome, though had not formed clearly independent groups. By day 14, the neuronal cells formed a tight cluster, with only slight differences seen between glutamatergic and GABAergic cell lines and minimal influence of *SLC39A8* genotype. Incorporating N-glycomics summary data from our detailed study of adult mouse cortex^22^ into the model showed that our differentiated neuronal cell types had close correlation to that of intact mouse brain tissue. Further, N-glycomics data from a human brain sample from the same study showed similar clustering in the model although further separated – however this single donor sample was from a commercial source with different processing. Day 14 wild-type endothelial cells clustered with a previously published N-glycome of primary human umbilical vein endothelial analyzed in a similar fashion (permethylation, MALDI-MS TOF) and timepoint (14 days in culture)^28^. In contrast, the N-glycome of day 14 endothelial cells lacking *SLC39A8* most closely associated with the group of day 4 induced cells. This finding suggests that, despite expressing a transcriptome consistent with mature endothelial cells (**Fig. 3**), differentiated endothelial cells lacking *SLC39A8* have impaired N-glycome maturation compared to wild-type cells.

**Figure 5.**
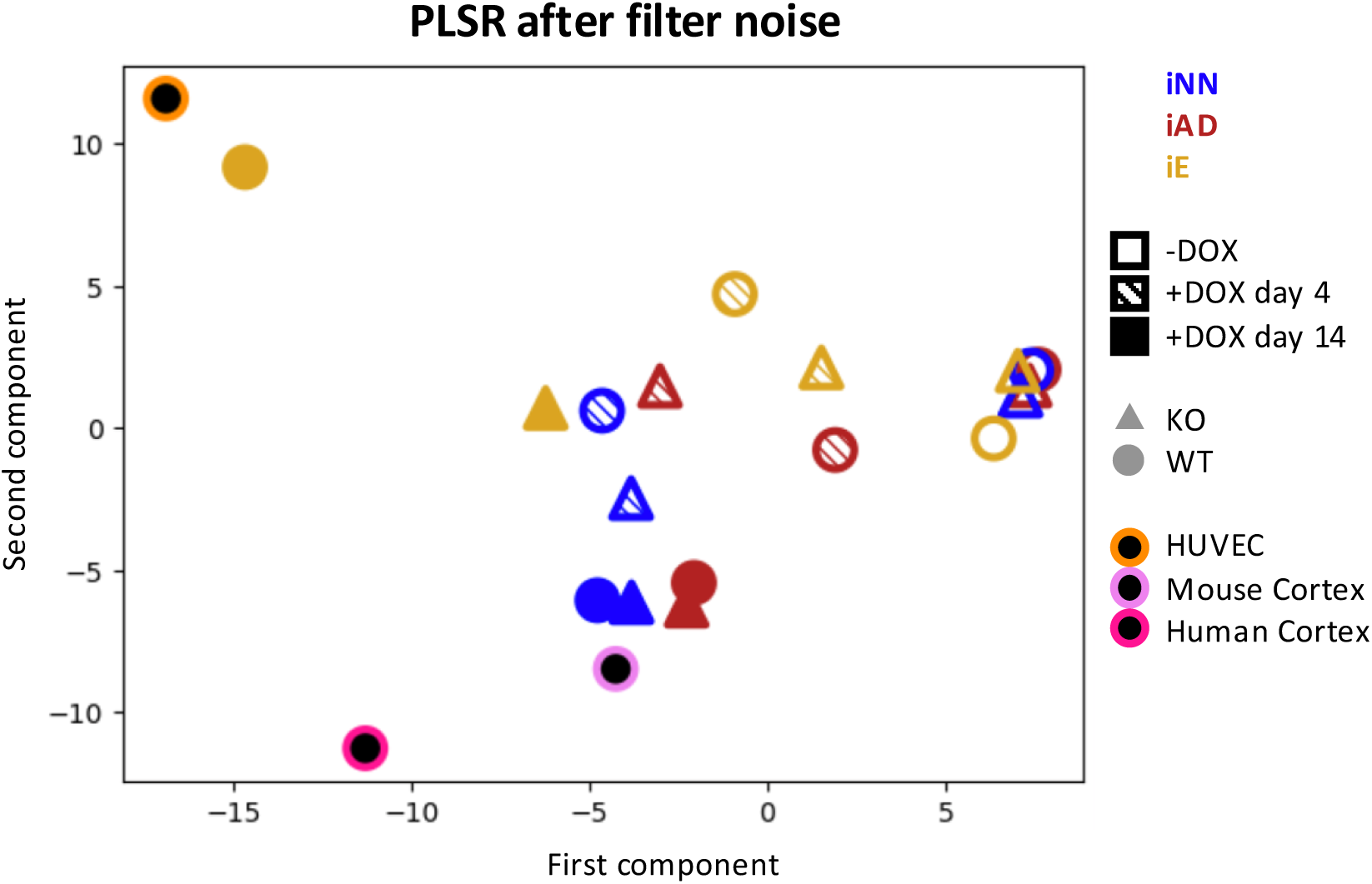
N-glycome profiles are unique between cell-type and resemble intact tissue. Plot of glycomic profiles from all cultures using Partial Least Squares Regression (PLSR) analysis, highlighting the tight clustering of each undifferentiated iPSCs line (-DOX, open shapes) compared to induced cell lines (+DOX, filled shapes, 4-day hash fill, 14-day solid fill) expressing distinct gene drivers for iNN-neurons (blue), iAD-neurons (red) and iE-endothelial cells (yellow). Data shown from two different genetic backgrounds, wild-type (circles) and *SLC39A8 -/*- (triangles) cells. Reference N-glycomics data from primary human endothelial cells (HUVECs) (black circle with orange halo) and mouse cortex (black circle with light pink halo show tight clustering with related 14-day wild-type samples. Human cortex (black circle with fuchsia halo) is located in similar location though further south-west compared to derived neurons and mouse cortex cluster.

### Combined transcriptome/analysis reveals unique secretory processing patterns of neurons through the Golgi

Given the distinct glycome profiles of iPSCs differentiating to neurons and endothelial cells, we assessed the expression and enzymatic products of the core Golgi glycosyltransferases involved in processing high mannose structures to generate complex N-glycans. Bottlenecks of glycan synthesis are more easily appreciated by arranging structures along their linear path of synthesis, akin to data from Bradberry and colleagues on fucosylation and synaptic vesicles^39^. Overlaying the subcellular localization and measured levels of each enzymatic transcripts provides an additional level of information on differences in the secretory pathway between cell lines (**Fig. 6A**). After leaving the ER, Man-9 is sequentially processed in the cis-Golgi by four distinct class-I α-mannosidases to generate Man-5. In the medial-Golgi, this structure is modified with a single GlcNAc by MGAT1, generating a glycan with one antenna (A1-Man-5). A1-Man-5 is then cleaved by two interchangeable class-II α-mannosidases, generating the pauci-mannose templates used by glycosyltransferases of the trans-Golgi to generate the complex N-glycan structures common to most intact tissues.

**Figure 6.**
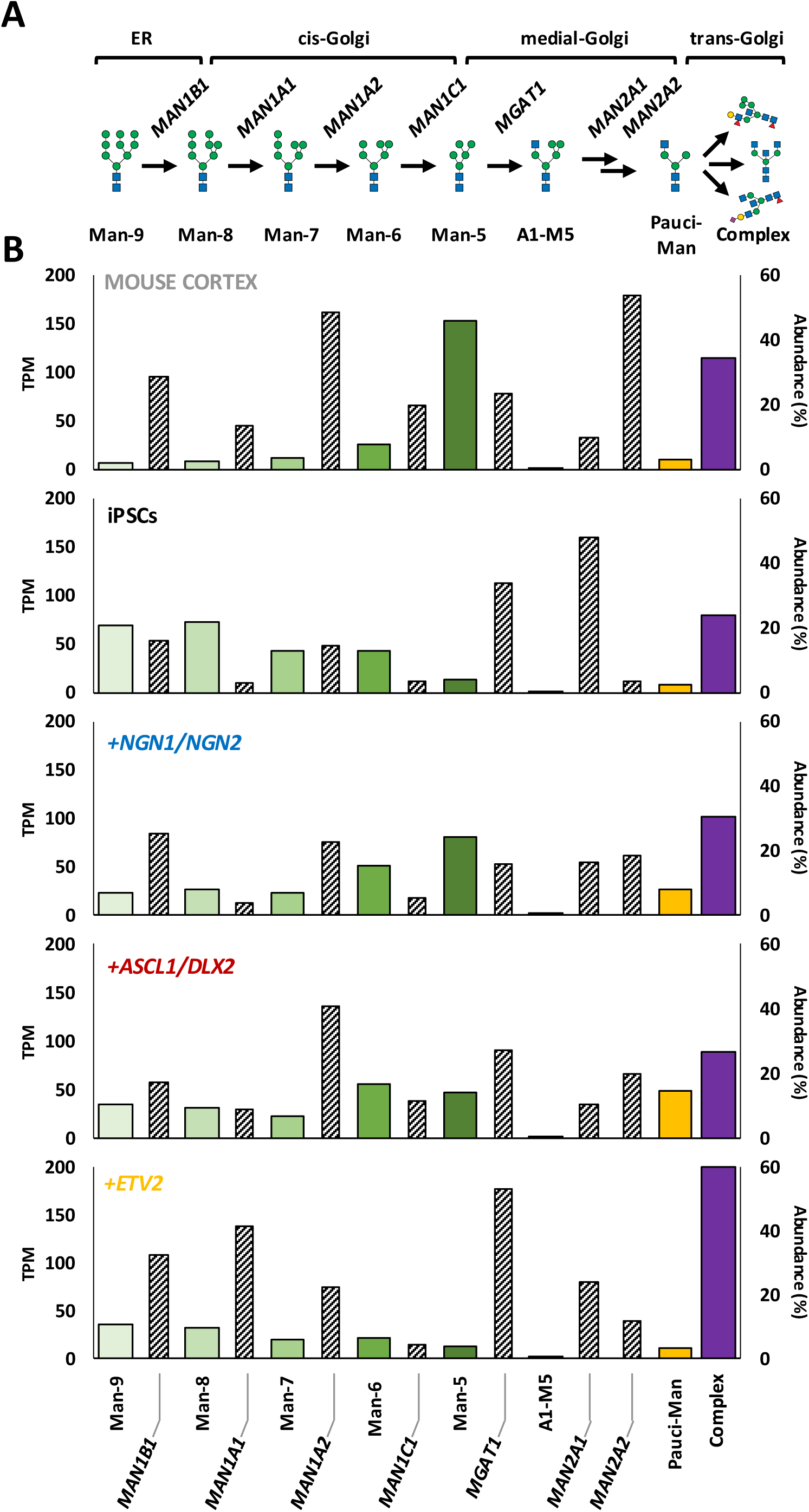
N-glycans of the cis-Golgi accumulate in neurons despite the presence of *MGAT1*. **A)** schematic of N-glycan synthesis from the ER through the Golgi, highlighting glycan types, glycogenes, and subcellular compartments. **B)** The relative abundance of individual N-glycans (%) from mouse cortex and differentiated neuronal cell types show a similar accumulation of the late cis-Golgi structures Man-6 and Man-5, while undifferentiated iPSCs and differentiated endothelial-like cells show an enrichment of structures from the ER and trans-Golgi, respectively. Glycogene transcript levels (diagonal stripes) for each step are shown as transcripts per million (TPM). Accumulation of the late cis-Golgi structures Man-6 and Man-5 in mouse cortex and differentiated neuronal cell types cannot be accounted for exclusively by an abundance of *MAN1C1* or a lack of *MGAT1*, as each express levels comparable to most tissues.

In our prior work^22^ and that of other groups using different analytic techniques^20,23,39^, the N-glycome of intact brain tissue shows an abundance of high mannose structures and a buildup of Man-5, which is consistently the most abundant N-glycan detected. In comparison, undifferentiated iPSCs show low levels of Man-5 and high levels of both early cis-Golgi structures (Man-9) and complex N-glycans having completed Golgi processing (**Fig. 6B**). In comparison, both iNN and iAD neurons exhibited a build-up of cis-Golgi products including Man-5 and Man-6, respectively, mirroring the intact brain N-glycome. The N-glycome of iE cells show lower relative levels of all high-mannose structures and a dramatic increase of complex N-glycans, similar to primary HUVECs^28^.

We and others have shown correlations between glyco-gene transcripts and their predicted products in brain and cell lines^22,36^. Thus, one potential explanation for the accumulation of Man-5 in neurons and the brain would be a lack of *MGAT1* expression, as is the case with Lec1 CHO cells^38^. However, RNAseq results from mouse cortex as well as each homogeneous culture confirmed comparable expression of *MGAT1* (**Fig. 6B**). Further, the expression of the four class-I alpha-mannosidases proximal and at least one of two interchangeable class-II α-mannosidases distal to *MGAT1* are expressed in all the cell lines to similar levels. These results further highlight the distinct N-glycome maturation process inherent to neuronal differentiation, which cannot be directly predicted by transcriptional analysis.

## Discussion

In this study, we used complementary methods on distinct cell-types differentiated from a shared hiPSC line to explore the cell-specific development of the brain N-glycome. We demonstrate that the unique profile of the brain is driven by neuronal cell types, both excitatory and inhibitory, and programmed into their differentiation. Further, we used this system to explore cell-specific effects of deleting the schizophrenia risk-gene *SLC39A8*, which, in contrast with most risk alleles, is expressed exclusively in endothelial cells.

The mechanisms regulating the distinct brain glycome and its function remain understudied^40^, particularly the connections and changes observed in disorders of the brain^41^. Given the complexity of the brain compared to other tissues, in which N-glycan complexity is relatively well distributed between high mannose, hybrid, and complex-type N-glycans in terms of abundance, it could be predicted that the N-glycome would perhaps be the most elaborate. However, the opposite is observed. Relatively few structures represent the majority of brain N-glycans, with a predominance of high-mannose, particularly Man-5, which is often considered simple as an intermediate precursor of more complex glycans with itself having unclear biological significance. Hanus and colleagues demonstrated the presence of high mannose structures on many critical proteins at the neuronal surface in culture^25^. They reported that blocking N-glycan processing beyond high mannose structures with kifunensine, deoxymannojirimycin, and swainsonine, all inhibitors of α-mannosidases, had no effect on dendritic arborization. In contrast, more proximal inhibition of N-glycan synthesis with tunicamycin, which blocks the initiation of N-glycan synthesis and high mannose generation on dolichol precursors, results in impaired dendritic arborization. Recent work from the same group showed that high mannose glycans are abundant at the neuronal surface in intact brains, and that changing their abundance alters the electrophysiologic properties of related circuits^42^. These studies provide support for a functional role of high mannose glycans at the surface of neurons and highlight the need for understanding the accumulation of these structures on a cellular level.

Several groups have investigated regional and developmental changes to the brain N-glycome across multiple mammal species. Lee and colleagues reported an in-depth study using LC-MS analyzed 9 different brain regions in adult mice, as well several timepoints in mouse and human prefrontal cortex^20^. They report a strong conservation of the overall N-glycome profile across each sample, with small but significant changes noted in less abundant structures including sialylated and fucosylated glycans^20^. Our group analyzed four brain regions (cortex, hippocampus, striatum, and cerebellum) in mice using MALDI-TOF MS and described a conserved N-glycome characterized by high mannose structures, as well as correlations between gene expression with glyco-transcripts involved in bisection and fucosylation^22^. Studies by Klarić and colleagues compared the same four brain regions in rat, macaque, chimpanzee, and humans at two time points and using combinations of chromatography and mass spectrometry, overlaid with gene expression data, and observed overall similar regional patterns^23^. In each study, subtle but statistically significant changes are observed between samples, and often used to draw conclusions regarding the function and evolution of the brain glycome. For example the change from ∼45.0% to ∼50.0% abundance of complex N-glycans from rat to human, or the reduction in α-2,3-linked sialic from ∼0.7% to ∼0.5% and a corresponding increase in α-2,6-linked sialic acid^23^. Analytical differences likely exist between studies, such as elution or ionization properties of different types of glycans in LC-MS vs MALDI. Despite this, an overlooked aspect of the N-glycome in every species, tissue, and time-point, is that high mannose glycans, and specifically Man-5, are always the most abundant structures in brain tissue. We show here that this build-up of Man-5 and its immediate precursor Man-6 is developmentally programmed into neuronal differentiation and should be considered the distinguishing feature of the mammalian brain N-glycome in comparison to other cell types and tissues, which commonly display much more complex N-glycans.

All eukaryotes share the same conserved core N-glycan processing steps and enzymes^17^, but different animals may differ in their overall diversity of N-glycans. For example, while both the drosophila and mammalian brain N-glycome are dominated by high mannose structures^43^, the peripheral tissues in drosophila are also primarily high-mannose structures, in contrast to the peripheral tissues of vertebrates, which display a diverse array of complex and modified N-glycans^44^. It is thought that the expansion in complexity of peripheral N-glycans of vertebrates, like other glycan classes, is driven by evolution, generating new carbohydrate structures at the cell surface for adaptive advantages such as managing the increased number of cell-cell interactions, regulating cell signaling, including interactions with microbes and pathogens^45–48^. The brain is separated from the periphery and pathogens by the blood-brain barrier, and thus may not require such complex glycans to identify and battle invading organisms. It is however the most complex organ in terms of cellular organization, cell-cell interactions, and cellular diversity. Thus, the restricted N-glycome of neurons and the brain compared to other cell types and tissues may result from not needing more complex N-glycans to function at some basal level, or, in contrast, that such strong constraint on the synthesis of complex N-glycans exists that relatively few are made. The conservation of the brain N-glycome across species, as well as the profound neurological symptoms which are hallmark of glycosylation disorders, suggests the latter is more likely.

A lack of MGAT1 expression would result in Man-5 accumulation, as is found in Lec1 cells^38^, though neurons express comparable levels compared to other cell types. *Mgat1*-null drosophila have decreased lifespan and abnormal brain development characterized by fused lobes^49^. Mice lacking *Mgat1* do not survive past mid-gestation (E9.5-E10.5), and show impaired neural tube formation^18^. The human *MGAT1* gene is under considerable genetic constraint^50^, and no known human disorders of MGAT1-deficiency have been reported. Targeted inhibition of *Mgat1* in the hippocampus of adult mice affected spine density and some electrophysiologic parameters including evoked potentials, but there was no report of histological neuronal death or phenotypes such as seizures in these animals^42^. As such, the role of high mannose and complex N-glycans during both development and normal function in the adult brain necessitate further study. Interestingly, all cells and tissues we have analyzed show a very low abundance of A1-Man-5 (< 0.7%), an essential intermediate structure for the synthesis of all complex N-glycans and the product of MGAT1. This suggests that the subsequent processing steps from A1-Man-5 by MAN2A1 and MAN2A2 to generate the pauci-mannose structures, upon which all complex N-glycans are built, is highly efficient when these enzymes can access their substrates.

An alternative explanation for the accumulation of Man-5 in neurons is the physical separation of maturing glycans in the secretory pathway from MGAT1. This is supported by several lines of evidence, including the existence of unique secretory processing and Golgi-independent trafficking of secretory vesicles in neurons^25,26^, as well as the weak correlation of glycosyltransferase expression levels with the observed N-glycome^23^. Further, glycome prediction tools based on glyco-gene expression levels, such as GlycoMaple, do not easily predict the buildup of Man-5 in brain tissue^36^. As such, although the transcripts for enzymes involved in generating complex N-glycans are expressed in neurons, their products many seldom interact with their substrates, or may do so only with a limited population or under specific conditions. Green and colleagues have described the existence of Golgi-satellites in neurons which show activity dependent activation of N-glycans^27^. A study by Stanley and colleagues identified an endogenous protein inhibitor of MGAT1, though this gene (*MGAT4D*) appears to only be expressed in the testis^51^. Spatial separation of different N-glycan species in neurons is also supported by enrichment of different glycan classes in certain subcellular structures^39^, as well as distinct patterns of N-glycan-binding lectins in the mammalian brain^52^. Lectins which bind high mannose glycans, such as ConA and GNL, show a diffuse pattern in the brain, while lectins for fucosylated and bisected N-glycans, specifically AAL and PHA-E, display enhanced signal in the synapse-rich molecular layer of the cerebellum.

Aside from the dramatic changes seen in congenital disorders of glycosylation and related mouse models, most glycome changes in common human diseases are modest in size^41^, and may relate more to downstream consequences of having the condition rather than playing a role in the etiology^53^. For example, our group compared the N-glycoproteome of 10 normal, 10 asymptomatic, and 10 symptomatic Alzheimer disease brain samples, and although some subtle differences were noted between groups, the overall profiles are strikingly similar^54^. An exception may be schizophrenia, where multiple glycosylation enzymes and related genes, including *SLC39A8*, are genetically associated with risk of developing the disorder^2^. In this study, we explored the cell-specific effect of *SLC39A8* deletion on the N-glycomes of neurons and endothelial cells. In contrast to the majority of schizophrenia risk genes, which display enriched expression in neurons^8^, *SLC39A8* is expressed exclusively in brain endothelial cells^16^. Akin to human plasma glycome studies^12,14^, we previously reported that mice harboring the schizophrenia risk variant A391T in *Slc39a8* showed changes in glycosylation of proteins expressed exclusively in neurons, suggesting a non-cell-autonomous effect of the schizophrenia risk variant on protein glycosylation. In line with those results, we find that *SLC39A8* deletion in neurons had minimal effect on their protein N-glycomes, whereas deletion in endothelial cells caused a dramatic reduction in the branching of complex N-glycans. Although some studies suggest that *SLC39A8* deletion or the A391T variant expressed in neuronal cell types can affect their function and contribute to schizophrenia risk^55^, the implications of these are unclear as *SLC39A8* is not endogenously expressed in these cell types^16^.

In summary, the results here emphasize both the unique N-glycome profile of neurons, which is programmed into their cell-specific differentiation, and the necessity of studying disease-relevant mutations in their appropriate cellular context. Future studies involving protein glycosylation in the brain should consider other less abundant brain cell types, including astrocytes, oligodendrocytes, and microglia, as well as other related pathways such as other glycosylation pathways, e.g., O-GalNAc glycosylation. Further, co-cultures and co-differentiation protocols will help understand the dynamic changes of the brain N-glycome and inform targeted treatments for abnormal protein glycosylation in the brain.

## Author Contributions

KK was involved in initial conceptualization of the project, generated all cell lines, performed differentiation experiments and cluster analyses for glycomic data, and wrote the manuscript

MN helped perform and optimize glycan assays and performed data analysis for glycomics experiments

MC performed analysis of RNAseq data

RIS designed and supervised RNAseq analysis

LW designed and constructed integration vectors and generated polyclonal hiPSC lines used in developing iE cells

JWS was involved in initial conceptualization of the project, experimental design, and data analysis

RDC was involved in initial conceptualization of the project, experimental design, data analysis, and oversaw all glycosylation analyses

RW was involved in initial conceptualization of the project, experimental design, data analysis, oversaw all cell generation and validation experiments, and wrote the manuscript

RGM was involved in initial conceptualization of the project, performed all glycomic experiments and data analyses, and wrote the manuscript

All authors contributed feedback and edits to the manuscript

## Acknowledgements

This work was supported by a foundation grant from the Stanley Center for Psychiatric Research at the Broad Institute of Harvard/MIT (awarded to RGM and RW) and R24GM137763 (awarded to RDC), P30DK040561 (awarded to RIS), and 1K08MH128712 (awarded to RGM).

## Methods

### Generation of Isogenic Cell lines

#### Cell Maintenance

Standard iPCS culture practices were used for all cell line maintenance. In brief, Matrigel (VWR Cat # 354277) coated plates were used with daily mTeSR+ (Stem Cell Technologies Cat #100-0276) media changes. Cells were aggregate or single-cell passaged, depending on the downstream application, with Gentle Cell Dissociation Reagent (Stem Cell Technologies Cat # 7174). Cells were gently thawed by rolling the cryovial between gloved hands until fully liquid and then placed directly into newly prepared plates with media. Rock inhibitor (StemCell Technologies Cat # 72304) was used in all media for thawing, passaging and in any case where colony size was significantly small, less than ∼10 cells. Cells were frozen down in dissociated suspensions of ∼1,000,000 cells per ml in mFreSR media (Stem Cell Technologies Cat # 05854). Mr. Frosty (ThermoFisher Scientific Cat # 5100-0001) was used to slowly cool cells during the freezing process. Cells were transferred to –140°C when fully frozen. All cell lines were generated from the same previously published human iPSC line, termed PGP1^29^. We confirmed that this cell line was derived from a male donor who was homozygous for the major allele (C) at rs13107325, which codes for alanine (A) at amino acid position 391 in *SLC39A8*.

#### Plasmids

Three base cell lines were initially created for this project, iNN (inducible circuit for glutamatergic neuron differentiation via *<u>N</u>GN1* and *<u>N</u>GN2* expression^30^), iAD (inducible circuit for GABAergic neuron differentiation via *<u>A</u>SCL1* and *<u>D</u>LX12*^31^) and iE (inducible circuit for endothelial cell differentiation via *ETV2*^32^). Addgene plasmids containing the Tetracycline-dependent promoter (TRE-tight) driving respective transcription factors (#61471 (NGN1/NGN2) or #97330 (DLX2) with #97329 (*ASCL1*)) were paired with an additional Lentivirus vector containing the core promoter for human Elongation Factor-1 α (hEF1a1) driving constitutive expression of reverse tetracycline-controlled transactivate (rtTA) to create the glutamatergic and GABAergic circuits respectively (Fig. 1A). The piggyBac integration system was used to create our ETV2 cell lines using a Golden Gate cloning strategy similar to previous work^56^. This modular cloning strategy was expanded to allow for construction of two transcriptional units (TUs) in a single piggyBac integration vector. For our PiggyBac transposon vector, the first TU consisted of TRE3G mediated *ETV2-2A-YFP*, the second TU used hEF1a to drive rtTA3, and the third TU applied hEF1 to drive the hygromycin resistance gene. To create cells lacking the expression of *SLC39A8* (SLC39A8-KO), a similar modular cloning strategy was used to create expression vectors for CRISPR/Cas9 and the U6 RNA polymerase III promoter (U6) driving guide RNA expression. Four guide RNAs were designed and initially tested on wild type PGP1. Only 1 combination gave successful modifications in the wild type PGP1, CHOPCHOP1 and CHOPCHOP3. Therefore, these two guide RNAs were used to make cuts that remove a 156 bp piece in both alleles in all three of our inducible cell lines (Supp. Fig. 1). Sequences for each guide RNA’s target sequence can be found in Supplementary material Figure 1b.

#### Lentiviral Production

Lentivirus was created using a 3^rd^ generation expression system through transient transfection of HEK293FT cells based on established protocols^57^. Fresh media was placed on wells 24 hr after transfection, and this media was harvested 48 hr after transfection. Virus-contained media was then filtered through a 0.45 μm polyethersulfone (PES) membrane. Virus was not frozen but rather used the same day of collection.

#### Lentiviral transduction

A Matrigel-coated 24-well plate of PGP1 cells at 50% confluency were incubated with filtered virus at 3 different concentrations, 1:4, 1:12 & 1:36 virus:media, in a total volume of 400 μl plus 10 μg/ml Polybrene mTeSR+ media with Rock for each well. Virus-containing media was allowed to sit on cells for 24 hours before being changed. The resulting polyclonal populations were expanded and underwent 2 passages before further analysis.

PiggyBac Integration: The piggyBac transposase mediated integration strategy was utilized to stably integrate our iE system^58^. In short, our *ETV2* PiggyBac transposon vector was co-transfected with a consultatively expressed PiggyBac transposase at the ratio of 4:1 for stable integration of our circuit. 48 hours after transfection, the stable integrated hiPSCs were selected with hygromycin for 14 days. The resulting polyclonal population was then expanded for sorting.

#### Single Cell Sorting, Expansion, and Selection

After stable integration of genetic circuits creating polyclonal populations, single cell sorts were performed to create monoclonal populations for each cell type. For iNN and iAD cell lines, sorting was gated based only on morphology to identify single cell populations, specifically three morphology gates were employed to isolate single cells based on forward and side scatter area, forward scatter width and height and side scatter width and height. Additionally, transiently transfected color controls were also used to back gate showing clear isolation of living single cells. For iE cells, sorting of single cells population was based on single cells displaying the top 10% of signal in the FITC channel to detect eYFP fluorescence, which should correlate with ETV2 expression based on the construct design harboring a 2A bicistronic peptide linker. Single cells from each line were sorted into 96-well Matrigel coated plates containing mTeSR+ media with Rock. After 1 day, sort media was replaced, and the single cells were allowed to expand for 5 days. After 5 days, media was removed and replaced with media without Rock, and cells left for an additional 3-5 days until the media color indicated decreased pH due to growth. Cells from one 96-well were moved to two wells of a 24-well plate. One of these wells was then further expanded into a 6-well plate, while DOX was added to the second well to assess for morphology changes over 2 days. If no morphological change was noted, both samples were discarded. For cells, that did show morphological positive samples, the morphological assay after DOX was repeated, and if again positive, the -DOX sample was selected and froze down at -80°C. Over 100 colonies were screened for each base cell line (iNN, iAD and iE), and of those that tested positive on the morphological screen (3-10%), one line for each base cell was selected and kept in culture for *SLC39A8*-knockout generation.

#### SLC39A8-Knockout (KO) Generation

Four guide RNA targeting human *SLC39A8* were designed using the CHOPCHOP web application^59^. Two of these four (CHOPCHOP1 & CHOPCHOP4) were designed to guide the system upstream of the Kozak sequence for *SLC39A8* and the remaining two (CHOPCHOP2 & CHOPCHOP3) were designed to guide the system downstream of the signaling sequence located in the first exon. All 4 logical combos were tested through transient transfection into undifferentiated PGP1 cell lines in a 24-well plate. Cells were grown for 2-3 days and then single cell sorted into Matrigel coated 96-well plates with Rock containing media. However, instead of the DOX morphology screen, the second split from the monoclonal sample was pelleted, and genomic DNA was extracted using QuickExtract (Lucigen Cat # QE09050), and a PCR over the region of interest was then conducted. During the initial screen only one of the four guide RNA pairs yielded a double positive KO (ChopChop1 with ChopChop3). Generation of KOs in the remaining cell lines iNN, iAD and iE used the top preforming pair of guide RNAs. One PCR product from a successful KO was gel extracted using Monarch DNA Extraction Kit (NEB Cat # T1020) and set for sequencing. The sequencing revealed the correct target region had been removed. One positive cell line from each cell population was chosen, resulting in 3 isogenic cell pairs with and without KO of *SLC39A8* (i.e. 6 cell lines total). Genomic PCR results from the final 6 cell lines confirmed successful deletion of the region of interest in SLC39A8 as described in results. All 6 cell lines further used in the study showed normal chromosomal reads as indicated by KaryoStat Assays conducted by Life Technologies (Catalog # A52849).

#### Transfection

Lipofectamine Stem Transfection Reagent (ThermoFisher Scientific cat# STEM00001) was used to transfect all plasmids in Matrigel coated 24-well plates as follows: 167 ng total DNA was prepared by using equal ratios of all plasmids prepped at 100 ng/μl involved in a particular transfection. Of note, no transfection marker, was used in order to reduce chances of off target effects. Optimem (ThermoFisher Scientific cat#31985062) was added to bring the total volume of each sample to 25 μl. A master mix of 1 μl Lipofectamine to 24 μl of Optimem was created, and 25 μl of this master mix was then added to each sample, creating 50 μl total volume in each sample. The complex was incubated at room temperature for 10 min before being added to the cells. Cells were then reverse transfected as single cell suspensions of 90,000 cells/ml and were added to Matrigel coated 24-well plates immediately before the transfection complexes were added to the cells to increase the surface area the transfection complexes could access on the cells. Plates were then incubated at 37°C overnight to allow cells to adhere, and the media was changed the following day.

#### DOX Induction

5 mg of Doxycycline Hyclate (DOX) (Sigma-Aldrich Cat# D9891) was resuspended in nuclease free water H_2_O to bring to make a 10 mg/ml stock, which was then aliquoted and stored at –20C. DOX was added to the media of newly seeded 6-well plates at a 5 μM concentration. Media was changed every 2-3 days for iNN and iAD cells, and daily for iE cells due to there more rapid proliferation.

#### Harvesting Cells

Undifferentiated iPSCs and iE cells were incubated at 37°C in Gentle Cell Dissociation Reagent (Stem Cell Technologies CAT#100-0485) for 5 minutes. Gentle Cell was aspirated and 1 ml PBS (VWR Cat#21-040-CV) was added, and cells were manually removed using a cell scraper, transfer to a 15 mL conical tube, and then pelleted at 300 G for 3 minutes. Supernatant PBS was carefully aspirated and pellets were flash frozen in liquid nitrogen. For iNN and iAD neurons, the Gentle Cell incubation was skipped as the neurons were easily dissociated from the plate. The media and cells were transferred to a 15 mL conical tube, and 1 mL was used to wash any additional cells off the plate and transferred to the conical tube, and the cells were pelleted at 300 G for 3 minutes. The supernatant was carefully removed, and pellets were flash frozen in liquid nitrogen. Of note, each culture and timepoint was grown with 6 technical replicates in a 6-well plate. Five technical replicates were pooled, pelleted and flash frozen for glycome analyses. One technical replicate was pelleted and frozen for RNA-seq analysis (note: we did not process the 4-day RNA-seq samples due to time constraints).

#### Immunofluorescence

iNN cells at 4 days +/- DOX were stained using standard protocols. In brief, live cells were gently washed with ice cold PBS 2x before being covered with 4% paraformaldehyde (Electron Microscopy Sciences Cat #157-4) for 20 min at room temperature for fixation. Fixed cells were then washed with PBS 3x and incubated with permeability/blocking buffer (10% donkey serum [Sigma Cat# D9663], 0. 2% Triton X-100 [MP Biomedicals cat #04807426] in PBS) for 1 hr at room temperature. Primary antibodies were incubated for 1hr at room temperature in the dark, washed with PBS 3x (5 min per wash), incubated with secondary antibody, when indicated for 1 hr at room temperature in the dark, washed with PBS 4x (5 min per wash), and then imaged using a Leica TCS SP5 II confocal microscope. The following antibody combinations/dilutions in permeability/blocking buffer were used: Anti-Sox2 conjugated to AlexaFluor647 (BD Biosciences Cat # 562139) at 1:200; Anti-Oct3/4 conjugated to AlexaFluor555 (BD Biosciences cat # 560306) at 1:100; Anti-Fox3/NeuN (BioLegend cat # 834502) at 1:1,000, donkey anti-mouse AlexaFluor546 (Life Technologies cat # A10036) at 1:2,000; anti-Tuj1 (Neuromics cat # CH23005) at 1:100, goat anti-chicken AlexaFlour633 (Life Technologies cat # A21103) at 1:2,000. Cultures were co-stained for either Oct4 and Sox2 or NeuN and Tuj1 together.

#### RNA isolation and Analysis

RNA from snap frozen cell pellets was extracted using the Monarch Total RNA Miniprep Kit (Cat # T2010S). RNAseq analysis was performed by the MGH NextGen Sequencing Core as previously described^15,22^. In brief RNA-seq libraries were prepared from total RNA using polyA selection followed by the NEBNext Ultra II Directional RNA Library Prep Kit protocol (New England Biolabs, E7760S). Sequencing was performed on Illumina HiSeq 2500 instrument resulting in approximately 30 million 50 bp reads per sample. Sequencing reads were mapped in a splice-aware fashion to the human reference transcriptome using STAR ^60^. Read counts over transcripts were calculated using HTSeq based on the Ensembl annotation for GRCh38 assembly and presented as Transcripts Per Million (TPM)^61^. The RNA-Seq statistical analysis (and related figures) was performed by using EdgeR package^62^.

#### MALDI-TOF N-Glycomic Analysis

All samples were processed as previously described^22^, with slight modifications as noted below. Cell pellets were lysed in 500 μL ice-cold buffer (50 mM Tris, 150 mM NaCl, 1.0% w/v Triton-X-100, pH 7.6) with protease inhibitor (Roche #46931320019), followed by brief dissociation using a hand-held motorized pestle (Kimble #749540) and 2 brief pulses of sonication for 10 seconds with a microtip (Qsonica Q700). Volume was adjusted to 1 mL with additional lysis buffer, and protein concentrations were measured using the Pierce BCA Protein Assay Kit (ThermoFisher Scientific #23255). Glycoproteins from 1 mg of protein lysate were then dialyzed, lyophilized, reduced, dialyzed, lyophilized, trypsinized, purified, eluted and lyophilized as previously described^22^. N-glycans were released from lyophilized glycopeptides after resuspension in 200 μL of 50 mM ammonium bicarbonate and incubated with 3 μL PNGase F (New England Biolabs, #P0704) at 37 °C for 4 h, then overnight (12–16 h) with an additional 3 μL of enzyme at 37 °C. Peptides were removed by a preconditioned C18 Sep-Pak columns (200 mg Waters, #WAT054945) glycans were eluted with 6 mL of 5% acetic acid, placed in a speed vacuum to remove organic solvents then and lyophilized prior to permethylation as previously described^22^. Permethylated glycans were resuspended in 25 μL of 75% methanol and spotted in a 1:1 ratio with DHB matrix on an MTP 384 polished steel target plate (Bruker Daltonics #8280781). MALDI-TOF MS N-glycomics data was acquired from a Bruker Ultraflex II instrument using FlexControl Software in the reflective positive mode with a mass/charge (m/z) range of 1,000–5,000 kD. Twenty independent captures (representing 1,000 shots each) were obtained from each sample and averaged to create the final combined spectra file. Data was exported in msd format using FlexAnalysis Software for subsequent annotation. For each set of cells, N-glycan masses were included if they had; 1- an isotopic appearance based on m/z, 2 - a mass that corresponds to a possible N-glycan composition, and 3 - the average signal to noise ratio was greater than 6 (S/N > 6). This resulted in the inclusion of 92, 80, 78, and 89 individual N-glycan masses for undifferentiated iPSCs, iNN, iAD, and iE cultures, respectively. The relative abundance of each glycan was calculated as the signal intensity for each isotopic peak divided by the summed signal intensity for all measured glycans within a spectrum. N-glycans were grouped into different categories based on shared components, such as monosaccharide composition, antennarity, etc., and the summed abundance of each category was compared. All glycan structures are presented according to the Symbol Nomenclature for Glycans (SNFG) guidelines^63^ and were drawn using GlycoGlyph^64^. A Partial Least Square Regression Analysis (PLSR) was performed using the normalized abundance from every peak that passed the criteria described above (isotopic, known composition, S/N >6) from any of the cell lines, resulting in 118 individual m/z values for comparison. If a peak was not originally annotated in the spectra for a cell line (not meeting the above-described criteria for the initial analysis) it was subsequently flagged in the other samples for the PLSR analysis such that there were no gaps in the peak data across samples. After normalization to relative abundance (signal intensity for each peak divided by the summed signal intensity for all measured peaks), the data was processed through a Savitzky-Golay filter to reduce noise before being regressed against the data corresponding day of DOX induction (0, 4 and 14). An optimal number of 4 components was found by minimizing the mean squared error of the cross validation between the predicted day and the actual day of DOX induction. The first 2 components were plotted against each other to assess clustering. Published data for permethylated N-glycans measured using similar techniques by MALDI-MS TOF from human cortex^22^, mouse cortex^22^, and primary endothelial cells^28^ were processed using the same filters and components and plotted for comparison.

## Data Availability

The data generated during this study are included in this published article and its supplementary information files, and available from the corresponding author on reasonable request. Raw MS glycomics and RNAseq data will be made available in public databases upon publication of this manuscript.

