## Supplemental Data for "The restricted N-glycome of neurons is programmed during differentiation"

This PDF file includes:
Figures S1 to S15
Table S1

A

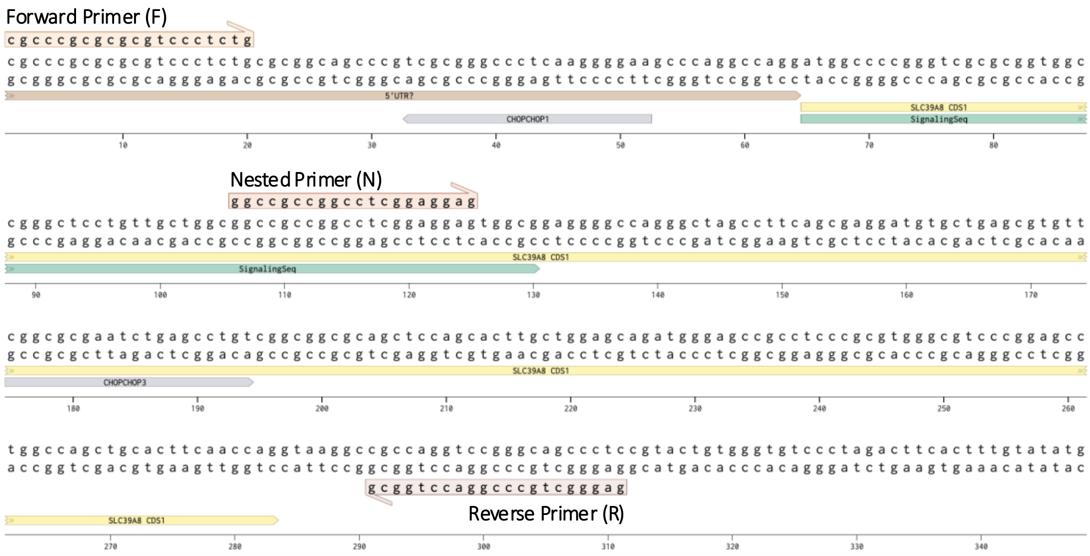


B

| Guide RNA | Sequence (5' → 3') |
|---|---|
| ChopChop1 | TTCCCCTTGAGGGCCCGCGA |
| ChopChop2 | CGCGAATCTGAGCCTGTCGG |
| ChopChop3 | CGGCGCGAATCTGAGCCTGT |
| ChopChop4 | TCTGCGCGGCAGCCCGTCGC |

C

| *SLC39A8* Genotype | F/R | N/R |
|---|---|---|
| +/+ (WT) | 311 | 206 |
| -/- (KO) | 149 | 0 |

D

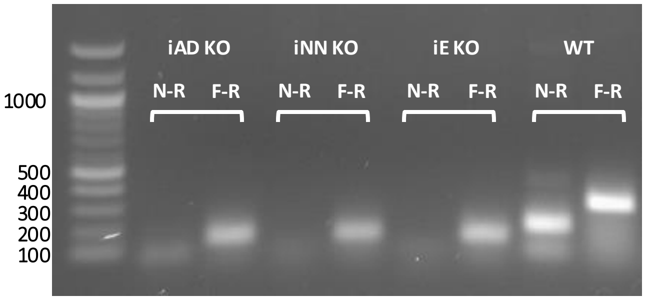


E

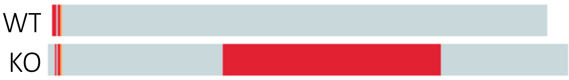


**Supp. Fig. 1. Genomic DNA Analysis of *SLC39A8* deletion using CRISPR/Cas9 in hiPSCs. (A)** Benchling schematic of guide RNA target sites and primer placement at the beginning of the *SLC39A8* coding sequence on chromosome 4. Primer sites are illustrated above and below DNA – Forward (F); Reverse (R); Nested (N). Cas9 sites (CHOPCHOP1/3, gray) are predicted to remove both the start codon and signaling sequence (green region) of *SLC39A8*, as well as the nested primer site. **(B)** Guide RNA sequences for deletion of SLC39A8. **(C)** PCR product size (bp) based on genotype and primer set. **(D)** Genomic PCR results confirm *SLC39A8* deletion for KO cell lines based on correct band size in in each case with the F-R primer set, and lack of a product with the N-R primer set. **(E)** Summary of sequencing results from gel extracted genomic PCR WT and KO (forward to reverse primer) bands. Grey represents regions of alignment between WT and KO, while red represents missing areas of the KO where there is no sequence to align, consistent with the expected region deleted by Cas9.

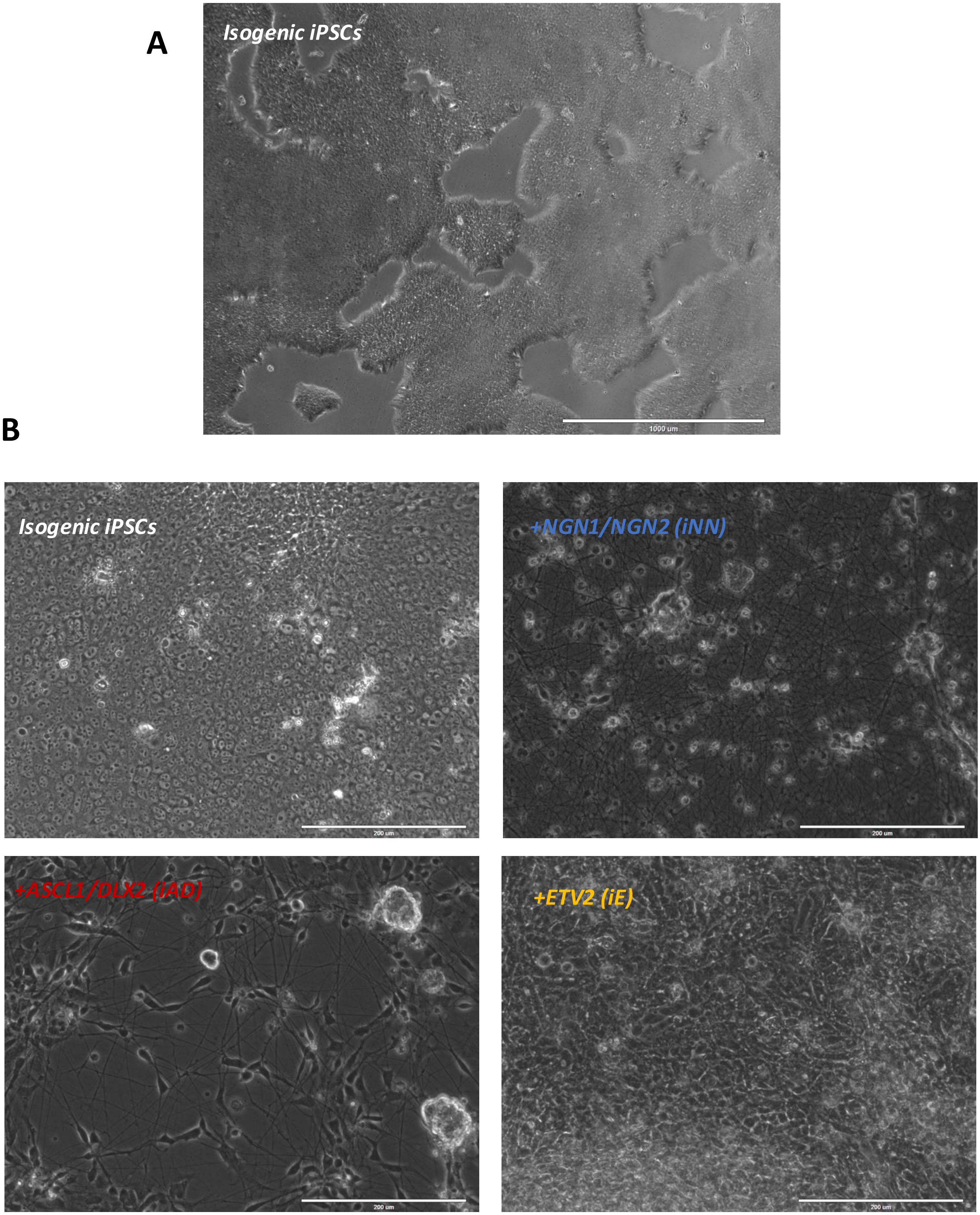


**Supplementary Figure 2. Light microscopy of differentiated iPSCs is consistent with cell-specific morphology. A)** Low magnification of undifferentiated iPSCs highlighting their sheet-like appearance in culture. Scale bar = 1 mm. **B)** High magnification of undifferentiated iPSCs and differentiated cell types after 14 days of +DOX induction. Scale bar = 200 µm.

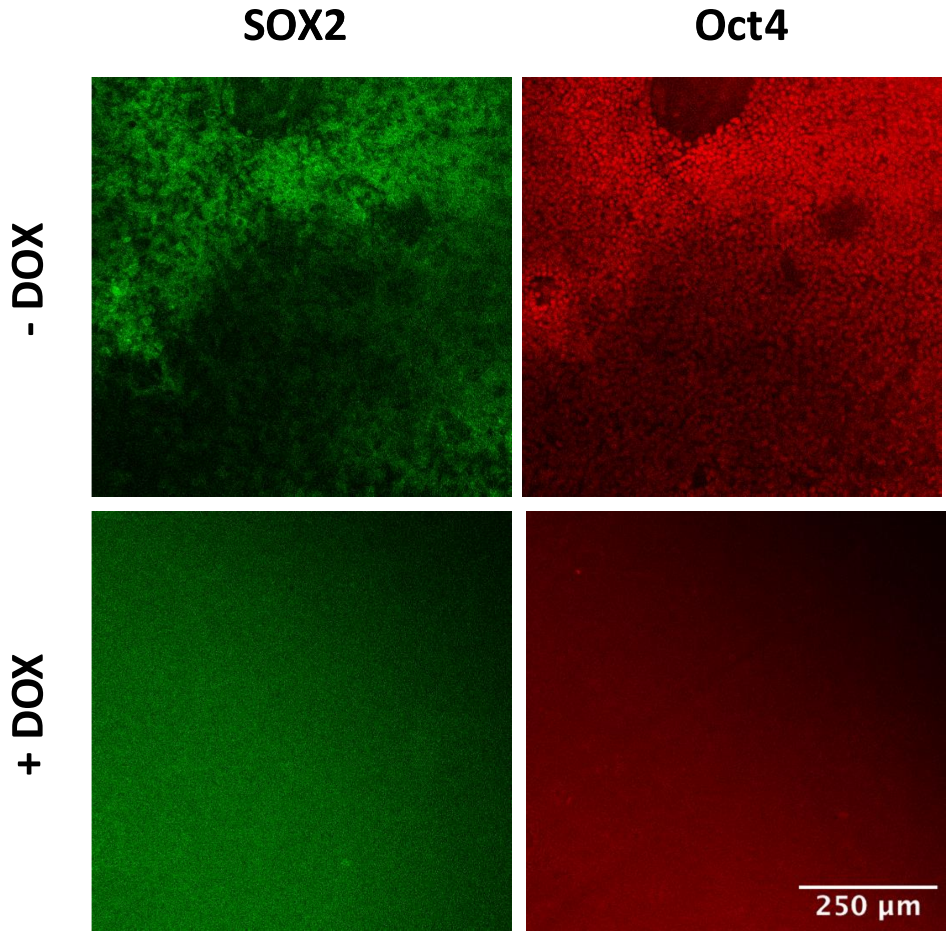


**Supplementary Figure 3. Immunofluorescence confirms the presence and absence of pluripotency markers in undifferentiated and differentiated NGN cell types, respectively.** Cultures of iNN iPSCs treated with and without doxycycline (DOX) to induce differentiation after 4 days were stained with stem cell pluripotency markers Sox2 and Oct4, which are present in the -DOX cultures and absent in + DOX cultures. Scale bars = 250 µM.

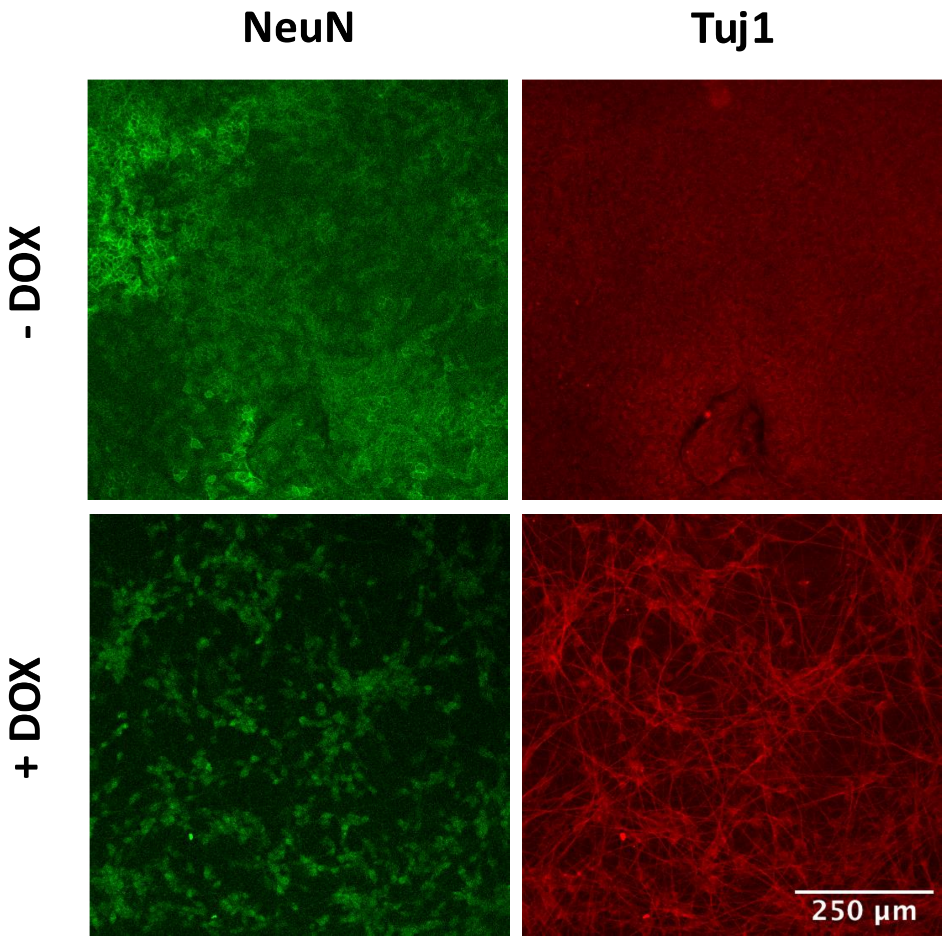


**Supplementary Figure 4. Immunofluorescence confirms the differentiation of neuronal (NGN) cell types in the presence of doxycycline.** Day 4 cultures of iNN iPSCs treated with and without doxycycline (DOX) to induce differentiation are stained with markers for neurons. NeuN (Fox-3) staining, a cytosolic protein which translocates to the nucleus in post-mitotic neurons, and positive Tuj1 staining, which binds the neuronal-specific class III β-tubulin, are consistent with the induced differentiation of neurons from iPSCs. Scale bars = 250 μM.

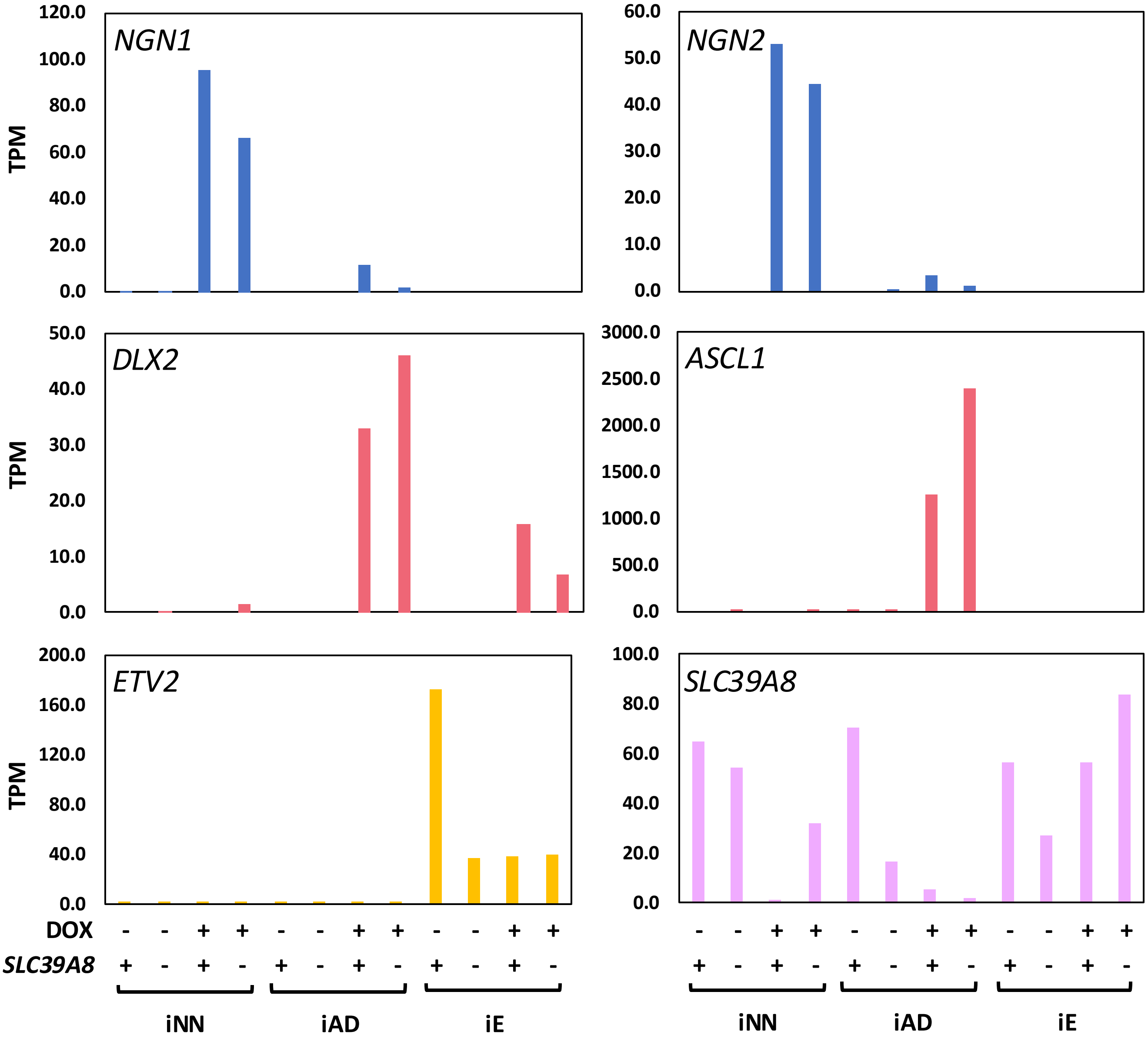


**Supplementary Figure 5. RNAseq profile of morphogen and *SLC39A8* expression across cell types.** Morphogens for neuronal induction were increased after DOX induction. ETV2 levels appeared unexpectedly high in the iE engineered cell lines based on RNAseq. *SLC39A8* levels decreased in mature neurons and appeared stable in uninduced and iE cells. +/- DOX induction and *SLC39A8* genotypes, including wild-type (*+/+*; **+**) and deleted (-/-, **-**) are indicated. Transcripts per million (TPM) shown on the y-axis.

A

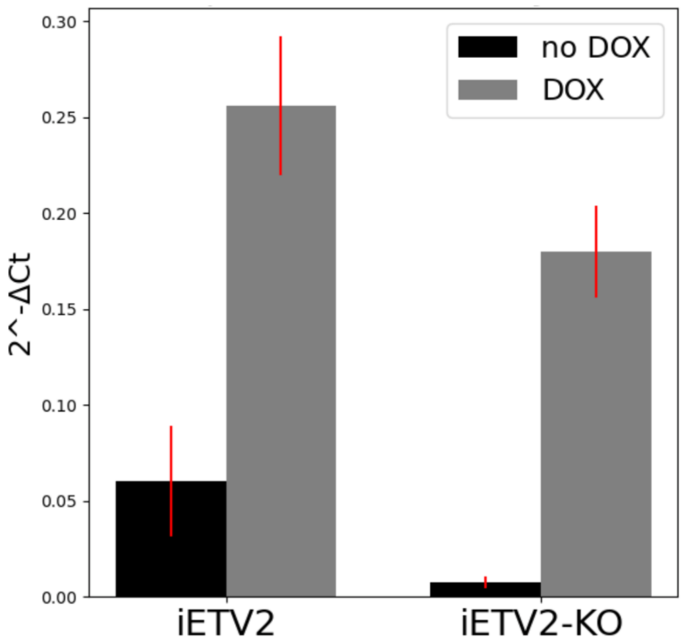


B

| Primer | Sequence | Efficiency |
|---|---|---|
| GAPDH_F | accacagtccatgccatcac | 134% |
| GAPDH_R | tccaccaccctgttgctgta | |
| ETV2_F | ggagcagagacacaaggaag | 103% |
| ETV2_R | cccttgctcaccataggtc | |

**Supplementary Figure. 6. qPCR confirms that endogenous *ETV2* expression is low in undifferentiated cells but increases dramatically with DOX in iE cells. A)** Delta Ct values calculated by comparing ETV2 levels to the reverence gene GAPDH after day 2 of +DOX. Experiment done in triplicate. Error bar are +/- 1 standard deviation calculated across all replicates. **B)** Primer sequences used to amplify GAPDH and ETV2 in qPCR experiment, with efficiency determined against DNA dilution standards.

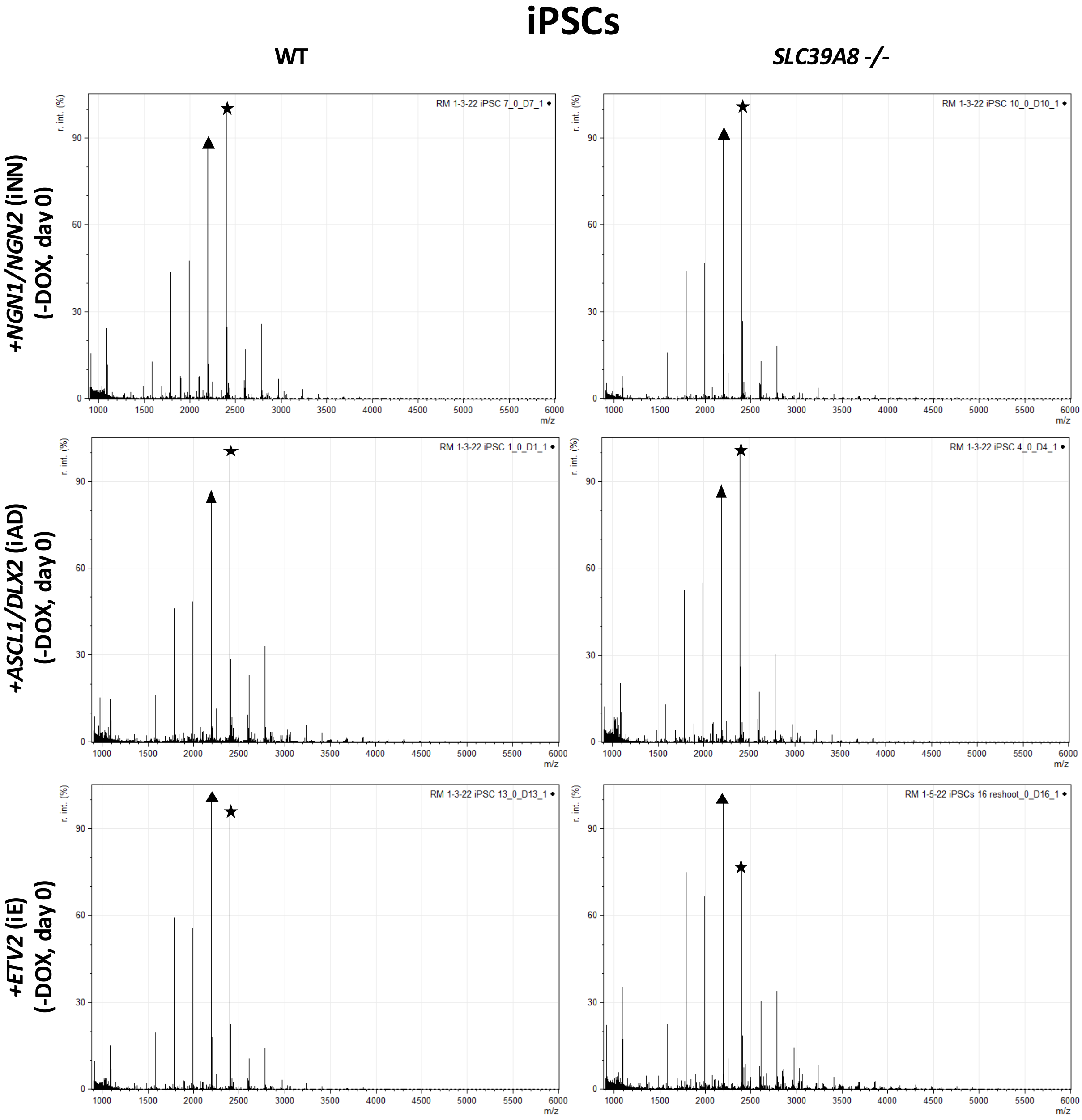


**Supplementary Figure 7. The N-glycome profile of undifferentiated iPSCs is characterized by an abundance of Man-9 and Man-8, and consistent across each cell line.** MALDI-MS TOF Analysis of permethylated N-glycans isolated from homogeneous cultures of each cell line (-DOX, day 0). In each culture Man-9 (★) and Man-8 (▲) are the two most abundant structures. Although the iE lines show a slightly greater abundance of Man-8 compared to Man-9, no glycan structures are significantly different across the undifferentiated cell lines.

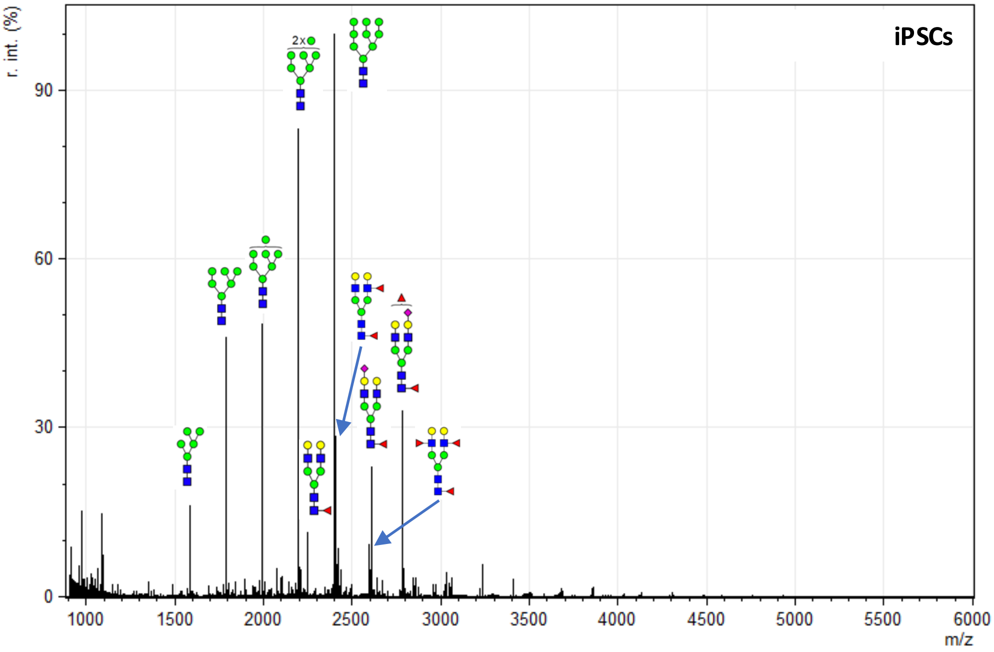


**Supplementary Figure 8. N-glycome profile of undifferentiated iPSCs.** Representative MALDI-MS TOF Analysis of permethylated N-glycans isolated from homogeneous cultures of undifferentiated (-DOX, day 0) iPSCs (AnD line). The corresponding structures for the 10 most abundant N-glycans, which make up over 82% of the total signal, are illustrated, including the larger high mannose species (Man-9, Man-8) and a few complex, highly fucosylated N-glycans.

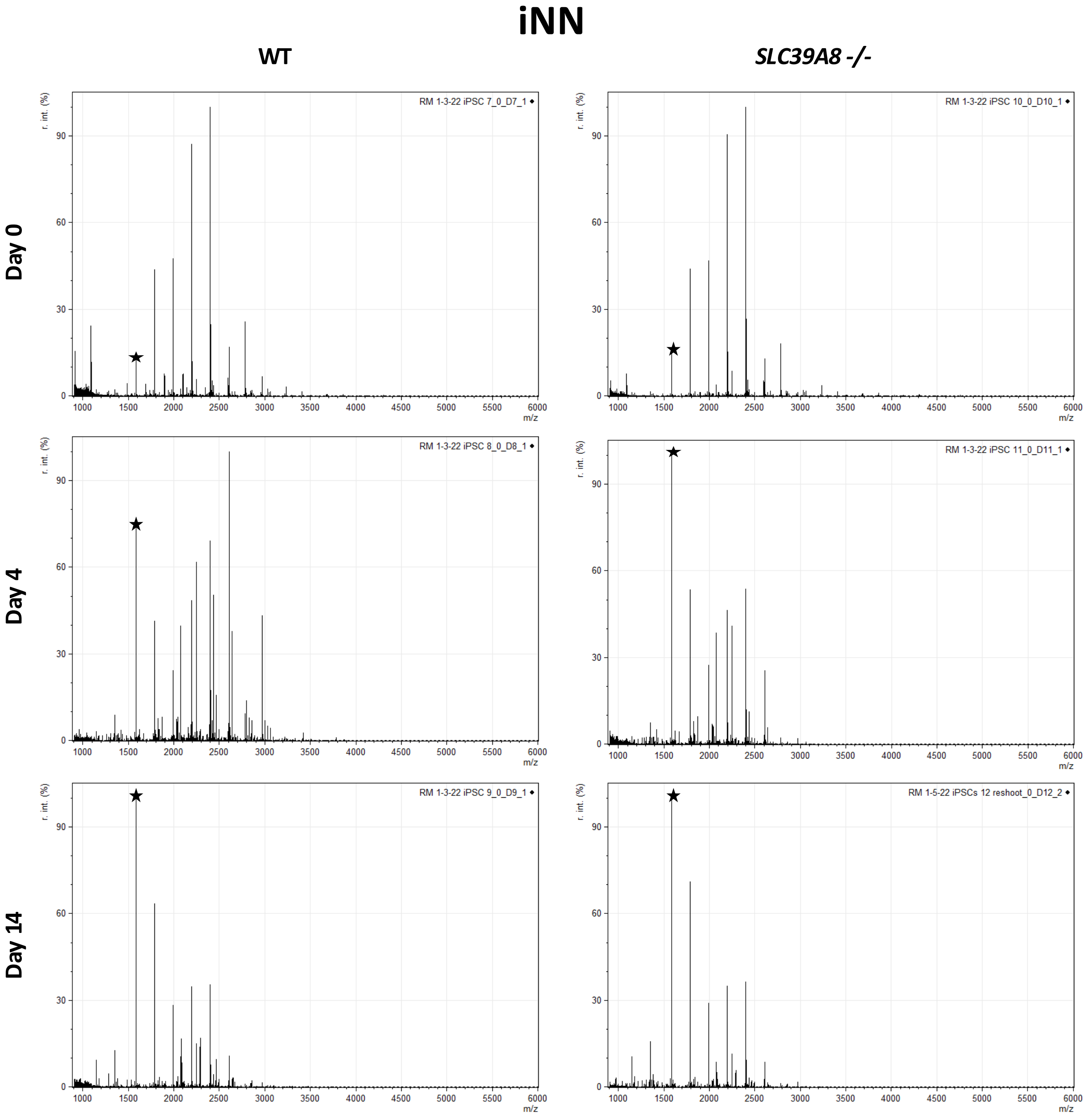


**Supplementary Figure 9. The N-glycome of iPSCs is dynamic during differentiation to glutamatergic neurons.** MALDI-MS TOF analysis of permethylated N-glycans isolated from homogeneous cultures at day 0 (-DOX), day 4 (+DOX), and day 14 (+DOX) from both wild-type (WT) and *SLC39A8* -/- iPSCs expressing *NGN1/NGN2* (iNN). In both genotypes, the predominant structures at day 14 are all high mannose, specifically Man-5 (★). This change is also evident at day 4 when the culture density and survival is lower than day 14.

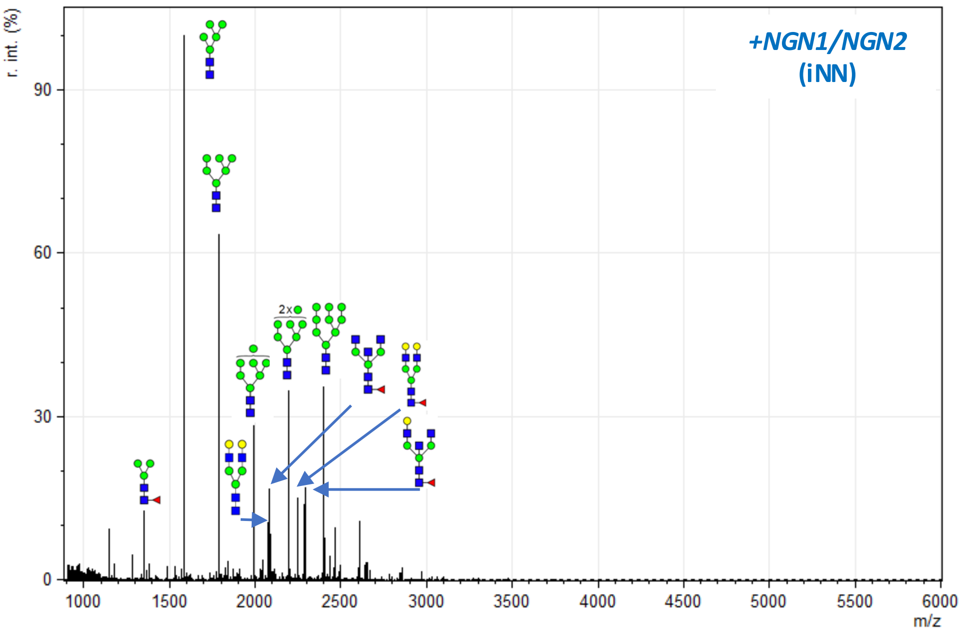


**Supplementary Figure 10. N-glycome profile glutamatergic neurons differentiated from iPSCs.** MALDI-MS TOF analysis of permethylated N-glycans isolated from homogeneous cultures of differentiated iPSCs (+DOX, day 14) expressing *NGN1/NGN2*, resulting in glutamatergic neurons (iNN). The corresponding structures for the 10 most abundant N-glycans, which make up over 77% of the total signal, are illustrated, including the smaller high mannose species (Man-5, Man-6) and a few complex N-glycans which include bisected structures and those with terminal galactose lacking sialic acid.

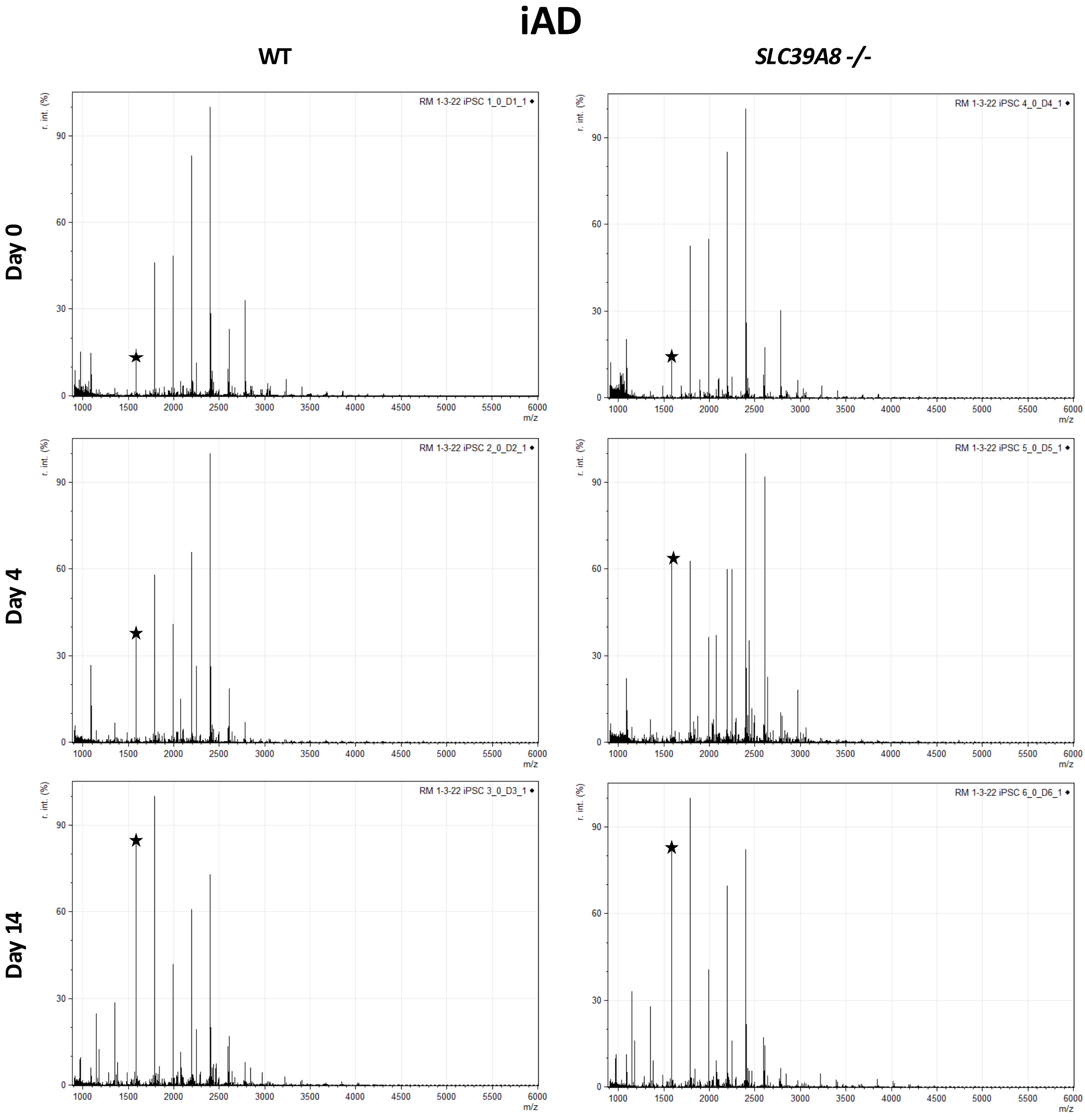


**Supplementary Figure 11. The N-glycome of iPSCs is dynamic during differentiation to GABAergic neurons.** MALDI-MS TOF analysis of permethylated N-glycans isolated from homogeneous cultures at day 0 (-DOX), day 4(+DOX), and day 14 (+DOX) from both WT and *SLC39A8-/-* iPSCs expressing *ASCL1/DLX2* (iAD). In both genotypes, the predominant structures at day 14 are all high mannose, including Man-5 (★). This change is also evident at day 4 when the culture density and survival is lower than day 14.

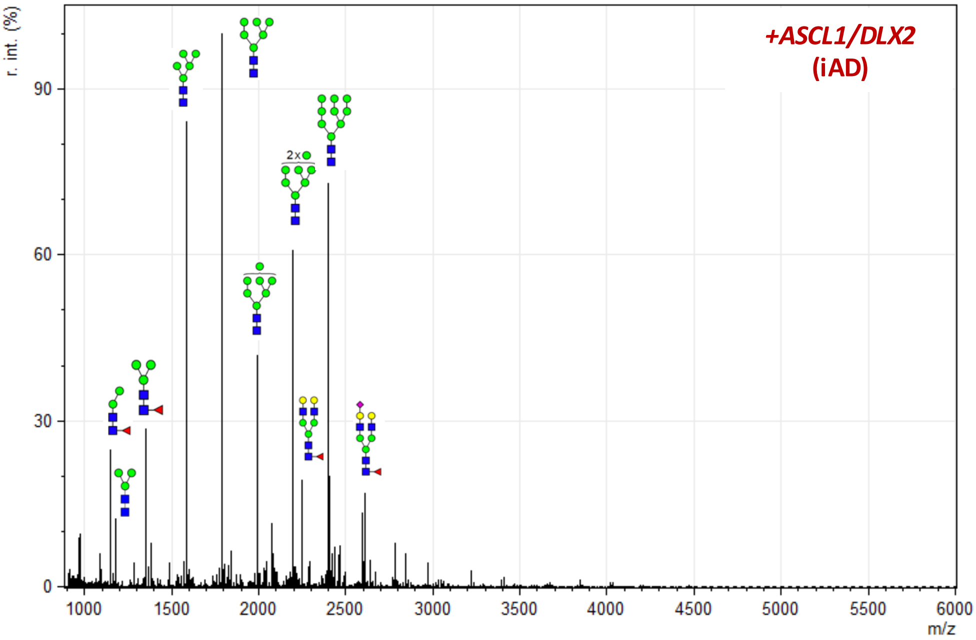


**Supplementary Figure 12. N-glycome profile GABAergic neurons differentiated from iPSCs.** MALDI-MS TOF analysis of permethylated N-glycans isolated from homogeneous cultures of differentiated iPSCs (+DOX, day 14) expressing *ASCL1/DLX2* (iAD), resulting in GABAergic neurons. The corresponding structures for the 10 most abundant N-glycans, which make up over 73% of the total signal, are illustrated, including the smaller high mannose species (Man-6, Man-5), pauci-mannose structures, and a few complex N-glycans.

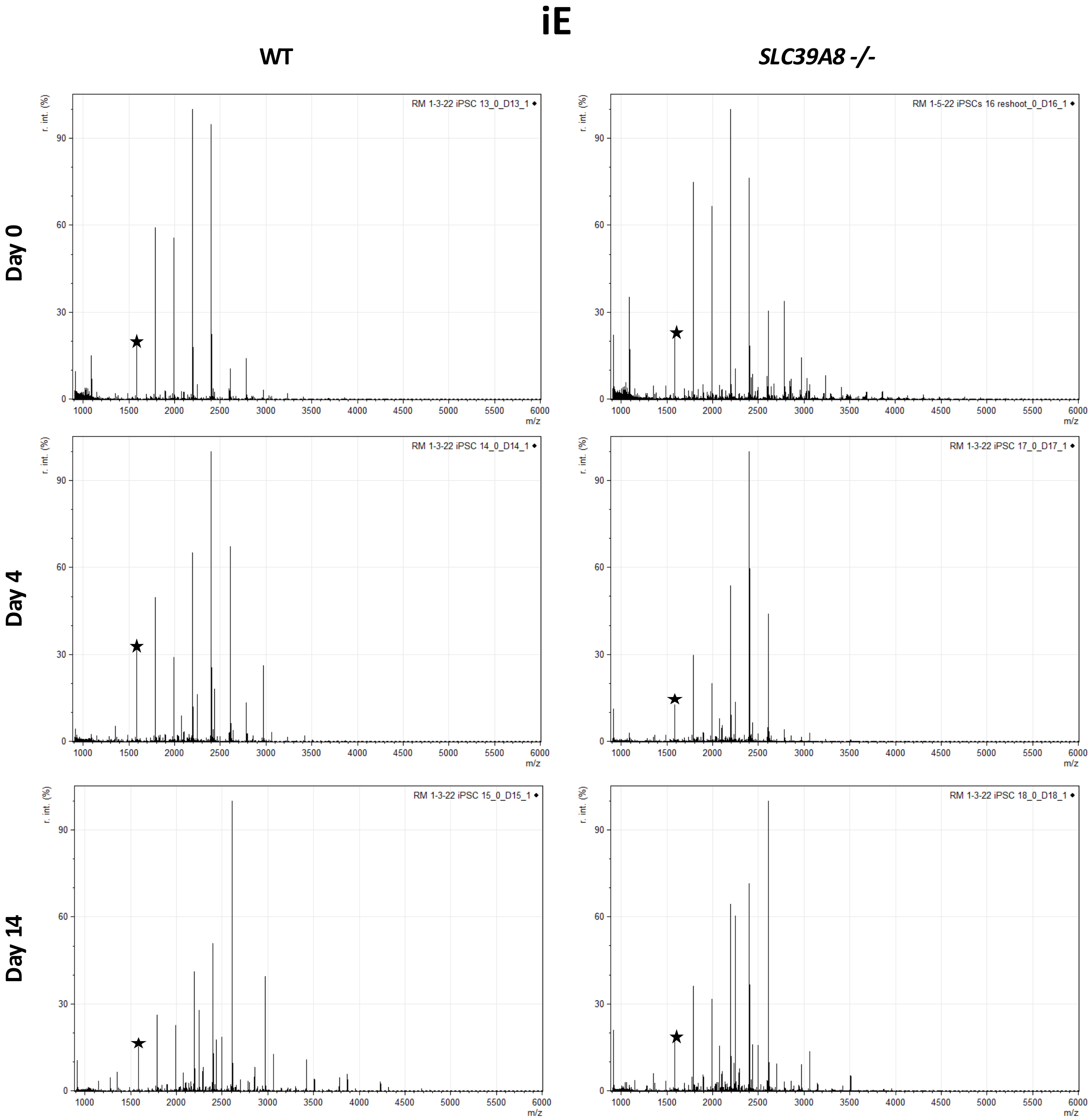


**Supplementary Figure 13. The N-glycome of iPSCs is dynamic during differentiation to endothelial cells.** MALDI-MS TOF analysis of permethylated N-glycans isolated from homogeneous cultures at day 0 (-DOX), day 4(+DOX), and day 14 (+DOX) from both WT and *SLC39A8-/-* iPSCs expressing *ETV2* (iE). In both genotypes, at day 14 the most abundant structure is the complex glycan FA2G2S1, with much less high mannose, including Man-5 (★). The increase in complex N-glycans is evident at day 4 in both cultures.

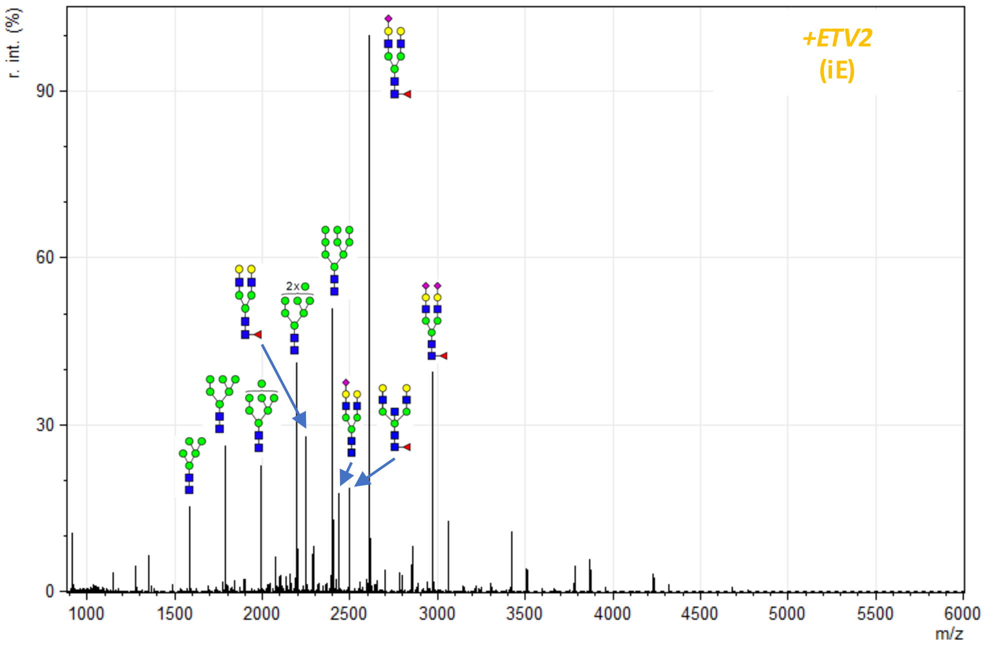


**Supplementary Figure 14. N-glycome profile endothelial cells differentiated from iPSCs.** MALDI-MS TOF analysis of permethylated N-glycans isolated from homogeneous cultures of differentiated iPSCs (+DOX, day 14) expressing *ETV2* (iE), resulting in endothelial cells. The corresponding structures for the 10 most abundant N-glycans, which make up over 75% of the total signal, are illustrated, including the complex N-glycans FA2G2S1 and FA2G2S2, and a few high mannose species.

A

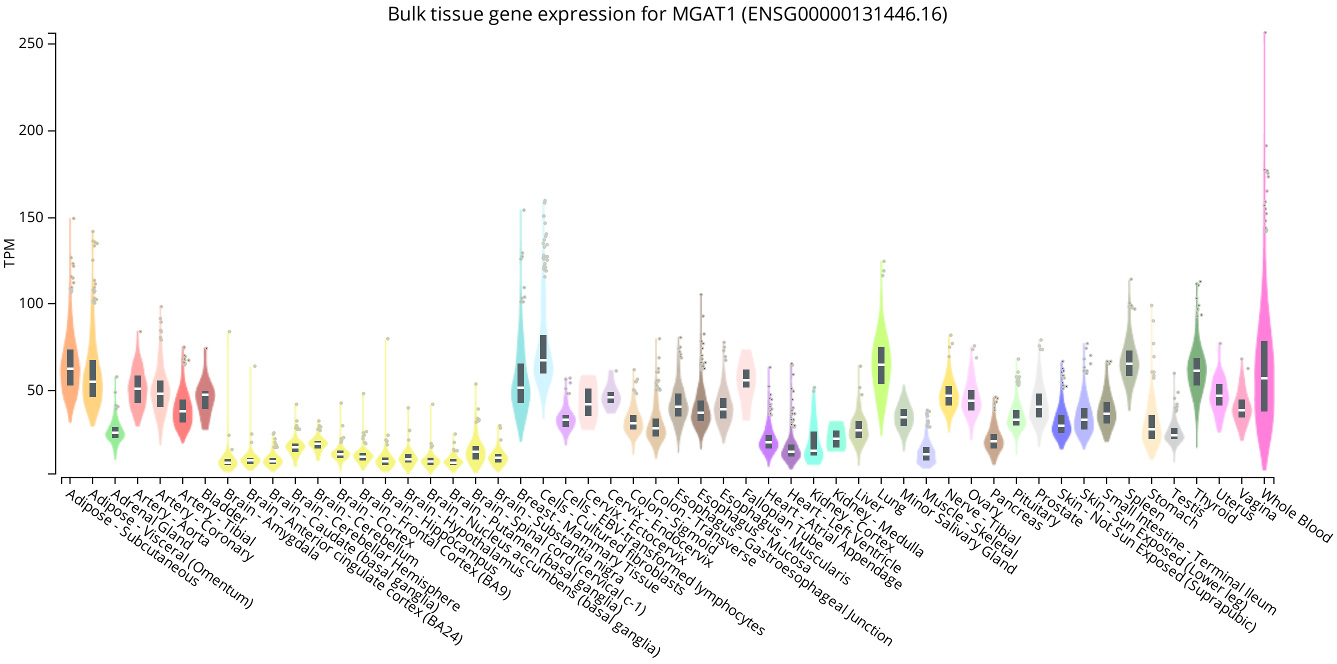


B

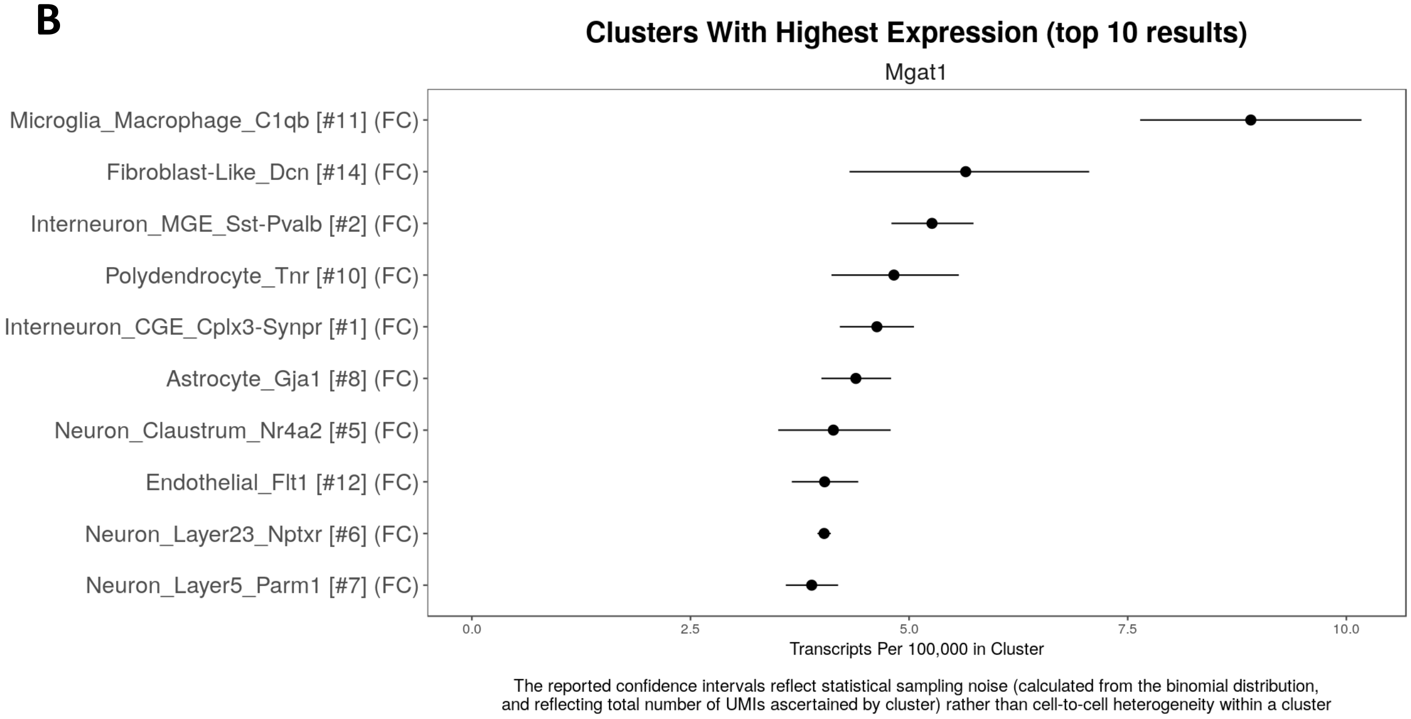


**Supplementary Figure 15.** ***MGAT1*** **expression across different tissues and single cell types of the brain.** A) Bulk RNAseq analysis of MGAT1 across each tissue type downloaded from the GTEx database (https://www.gtexportal.org). *MGAT1* shows broad expression but decreased levels in the brain, consistent with the restricted expression of glycogenes in the brain compared to other tissues. B) Single cell expression data of *Mgat1* in mouse cortex downloaded from the DropViz database (http://www.dropviz.org). *Mgat1* is detected at low levels in each cell type measured.

| | iPSC | iE |
|---|---|---|
| **ENDOTHELIAL MARKERS** | | |
| ***ERG*** | 1 | 77 |
| ***FLI1*** | 1 | 91 |
| ***TAL1*** | 0 | 25 |
| ***CDH5*** | 0 | 520 |
| ***CLDN5*** | 1 | 259 |
| ***VWF*** | 3 | 181 |
| ***FLT1*** | 102 | 482 |
| ***TIE1*** | 13 | 323 |
| **EPITHELIAL MARKERS** | | |
| ***MUC1*** | 4 | 1 |
| ***EPCAM*** | 390 | 246 |
| ***CDH3*** | 423 | 70 |
| ***CDH1*** | 341 | 71 |
| **Manganese Transporters** | | |
| ***SLC39A8*** | 27 | 84 |
| ***SLC30A10*** | 3 | 6 |
| ***SLC39A14*** | 463 | 208 |

**Supplementary Table 1. Transcriptomic profiling shows that ETV2 induction generates brain-like endothelial cells.** Expression levels for markers of brain-like endothelial cells are increased following ETV2 induction, while markers of epithelial cells are generally low or decreased. Levels of several common manganese transporter are shown. Data for each gene is shown as transcripts per million (TPM).
